# CETP alternative splicing variation impacts human traits

**DOI:** 10.64898/2026.03.16.712143

**Authors:** Isabel Gamache, Marc-André Legault, Jean-Christophe Grenier, Éric Rhéaume, Jean-Claude Tardif, Marie-Pierre Dubé, Julie G Hussin

## Abstract

The cholesteryl ester transfer protein (CETP) is an important protein in reverse cholesterol transport and has been identified as a significant factor associated with cardiovascular disease (CVD), making it a widely studied pharmaceutical target. Three protein-coding isoforms of *CETP* exist, distinguished by the alternative splicing of one exon each. The isoform primarily responsible for cholesterol-related functions in the plasma is well studied, but specific functions of each isoform remain poorly understood. In this study, we demonstrate the significance of considering CETP’s isoforms in analyses of human traits. Using bulk RNA-seq data from multiple tissues, we characterized the expression patterns and genetic regulation determinants of *CETP* transcripts. Leveraging publicly available GWAS summary statistics, we conducted multivariable Mendelian Randomisation (MVMR) to estimate the impact of variation in isoform proportions on phenotypes, highlighting the importance of *CETP’s* isoforms in pituitary and thyroid glands. Furthermore, we uncovered tissue-specific associations between *CETP’s* isoforms and CVD-associated phenotypes. Additionally, we observed that the epistatic interaction previously reported between *CETP* and *ADCY9*, a gene implicated in modulating a CETP modulator’s response, may be mediated through the regulation of alternative splicing of exon 9. Our results underscore the importance of a comprehensive understanding of *CETP’s* isoforms, which can significantly impact both fundamental and clinical research efforts.

## Introduction

The cholesteryl ester transfer protein (CETP) mediates the exchange of cholesterol esters (CE) and triglycerides (TG) between high-density lipoproteins (HDL) and Apo-B containing lipoproteins(1,2). It is expressed in a wide variety of tissues (3–6), and plasma CETP is mainly produced by liver and adipocytes (6). Many factors can modulate its production, including dietary cholesterol(7–9), thyroid stimulating hormone (TSH)(10), hyperinsulinemia (11), obesity (12) and acute infection with Epstein-Barr virus (EBV)(13). CETP’s function in plasma is mostly linked to the modulation of lipoprotein metabolism, but intracellular functions have also been reported, notably linked with lipid homeostasis, such as lipid transport from the endoplasmic reticulum (ER) to lipid droplets, TG biosynthesis, lipid storage, or its cholesterol content in the membrane (1,2,14–20). CETP’s impact on lipid concentration in different lipoprotein particles has led to the hypothesis that its pharmacological inhibition leading to increased HDL-cholesterol (HDL-C) concentration may be beneficial in treating cardiovascular diseases (CVD) (21–24), resulting in the development of several CETP inhibitors (25–29). These inhibitors act on lipoprotein-associated lipid profile in plasma and can also act within cells (30–34). Other phenotypes have also been associated with this gene, such as response to sepsis (35–40).

While previous expression quantitative trait loci (eQTL) studies examined the impact of genetic variants on global *CETP* expression levels, there is a lack of research exploring *CETP*’s expression regulation at the mRNA isoform-level. CETP has three reported protein-coding transcripts: the full-length transcript (Ensembl CETP-201 transcript), the exon 9-spliced out transcript (CETP-202), and a transcript featuring an alternative first exon (CETP-203). A majority of studies have focused on CETP-201, as it is the predominant form that is secreted from most cells and exerts plasma activities (6). Meanwhile, the protein derived from CETP-202, which is not secreted, functions as an inhibitor, binding to CETP-201 protein and impeding its secretion (4,41–44). Notably, to our knowledge, there is currently no clear knowledge about CETP-203’s function, which is nevertheless reported as a coding protein in Ensembl (45).

Alternative splicing refers to the process by which exons from the same gene are joined in various combinations during mRNA maturation leading to distinct isoforms, which can exhibit different functions and impact on phenotypic traits (46–50). Genomic regions surrounding and within an alternative exon are important for the regulation of the splicing events since they can impact the recognition of splicing binding factors. In CETP, exon 9 is spliced out in the *CETP-202* isoform and the exonic mutation rs5883 within this exon influences the recognition of a splicing factor, thereby modulating the occurrence of this splicing event (51–53). This mutation was also found to be linked to HDL-C level and coronary artery disease (CAD) in a sex-specific manner (51). Interestingly, an intronic mutation located near exon 9, rs158477, in strong linkage disequilibrium (LD) with rs5883 (in the CEU population from 1000 Genomes project (54)), is involved in an epistatic interaction with *ADCY9* impacting *CETP* expression levels (55). This epistatic interaction appears to have been under sex-specific selection in the Peruvian population, with an over-representation of haplotypes involving rs158477 in *CETP* and rs1967309 in *ADCY9* that differ between males and females. Rs1967309 genotype located at approximatively 50 MB from the *CETP* gene determines the CVD benefits in response to the CETP modulator dalcetrapib (56).

In this study, we characterized the genetic regulation of expression and splicing for each *CETP* isoform using data from the Genotype-Tissue Expression (GTEx) project. Furthermore, we demonstrate that changes in the proportion of *CETP*’s isoforms have a significant impact on various phenotypes using Mendelian Randomisation (MR) analysis based on publicly available summary statistics. Moreover, our findings indicate that changes in the proportion of *CETP* isoforms are associated with CAD in a tissue-specific manner. Additionally, in line with previous results on the evolutionary link between *CETP* and *ADCY9* genes (55), our study shows that the interaction between these genes has a significant impact on alternative splicing of exon 9. Lastly, variations in the proportion of *CETP* isoforms may indicate important functions of CETP within the pituitary or thyroid gland, highlighting the need for further investigation into the role of these isoforms in physiological processes. Generally, our study highlights the crucial importance of studying the individual isoforms of CETP.

## Results

### Tissue-Specific Genetic Regulation of *CETP* Isoforms Reveals Distinct Regulatory Patterns

*CETP* regulatory region has been previously described in the literature, with tissue-specific eQTLs located in the upstream region of the gene (8). Its expression has been detected in many tissues, but is typically mostly found in adipose, breast, spleen and liver (53). Since most of the studies that examine *CETP* expression only evaluate overall gene-level expression, hence ignoring isoform-level patterns, our goal is to study differences in genetic regulation in an isoform-specific way to determine their specificities. Using GTEx RNA-seq data, we confirmed that the most expressed isoform is the full-length transcript, *CETP-201*, whereas the alternative exon 9 transcript, *CETP-202,* and the isoform with an alternative starting exon 1, *CETP-203*, are less expressed in all tissues except in Lymphoblastoid Cell Lines (LCL) (Supplementary text, Supplementary Figures 1 and 2). To study the genetic regulation of each of the *CETP* isoforms and compare them to gene-level *CETP* regulation, we identified *CETP* eQTLs, within a 20 kb region surrounding the locus (Methods). As previously reported in the literature (8), significant gene-level eQTLs are found upstream of the gene, in the LD block B-3 (Figure 1), and show heterogeneity across tissues, confirming a tissue-specific regulation (Methods, Supplementary text, Supplementary Figure 3). The impact of the most statistically significant eQTL, rs56156922 (Figure 1), strongly varies across tissues (Supplementary Figure 3), with the small intestine showing the strongest effect, potentially linked to its role in cholesterol absorption (8). Notably, in the testis and ovary, rs56156922 exhibits opposite effects on *CETP* expression compared to other tissues (Supplementary text), suggesting a potential involvement of these organs in sex-specific traits associated with CETP activity, since eQTLs exhibiting opposite effects between tissues have been suggested to play a role in the development of complex trait (57).

**Figure 1.**
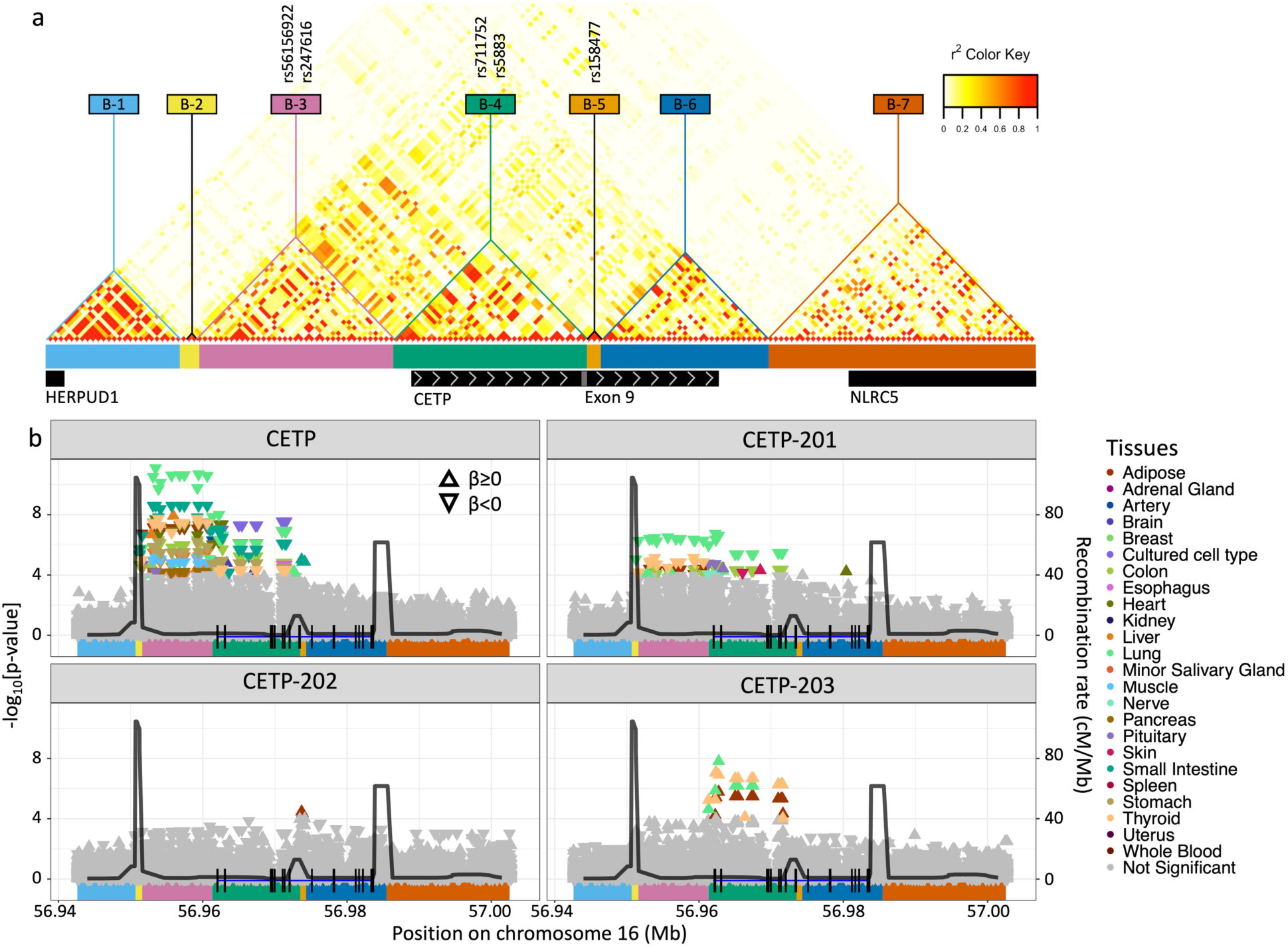
Gene Structure and Genetic Regulation of *CETP* Transcript Expression. (a) Linkage Disequilibrium (LD) blocks within the *CETP* locus, with surrounding genes identified below and grey arrows indicating 5’ to 3’. Blocks were estimated in GTEx participants from European descent using BigLD of gpart package with CLQ cut at 0.3 for SNPs having a MAF above 5%. IDs of SNPs of interest in this study are localized within their respective LD blocks. Position of the exon 9 within CETP is represented by a grey box. (b) cis-eQTLs of gene-level *CETP* expression (CETP) and its transcripts (CETP-201, CETP-202, CETP-203) for 49 tissues of GTEx, grouped by 24 tissue labels. LD blocks are represented under each plot using the color code from (a). SNPs below the threshold (p-value<0.05/[49 tissues x 7 LD blocks]) are colored. The CETP exons are represented in each plot, specific to each isoform. Black lines represent the recombination rate in CEU of 1000G.

To investigate isoform-specific regulation, we compared the estimated effects of significant eQTLs of each isoform to those of gene-level *CETP* (Methods). The genetic regulation of gene-level *CETP* is not significantly different from that of *CETP-201* and *CETP-202* (Supplementary Figure 4), with *CETP-201* showing significant association within the same LD block and displaying tissue-specificity (Supplementary text, Supplementary Figure 3). *CETP-202* does not exhibit distinct significant eQTLs in the *CETP* locus, suggesting that both isoforms are regulated by the same genetic region. In contrast, distinct significant eQTLs are found for *CETP-203,* located in LD block B-4 (Figure 1), which showed significant differences in effect sizes compared to gene-level, *CETP-201* and *CETP-202* eQTLs (Supplementary Figure 4) in tissues where *CETP* is the most expressed, although there was no clear evidence of tissue-specific effects for *CETP-203* (Supplementary text). These results indicate that *CETP-203* is differently regulated from the other two isoforms and points towards a novel putative regulatory region not captured by gene-level eQTL analysis.

### Genetic Regulation of CETP Isoforms Revealed through Alternative Splicing Analysis

Analyzing *CETP* eQTLs at the isoform level did not result in distinguishing the specific regulatory mechanisms of the main isoforms, *CETP-201* and *CETP-202*, hindering their comprehensive characterization. However, an approach based on alternative splicing (AS) can provide a better understanding of their distinct genetic regulations. *CETP* isoforms *CETP-202* and *CETP-203* arise from distinct splicing events, specifically, the alternative splicing of exon 9 (AS9) (Figure 2a, Junction 2) and alternative exon 1 (AS1) (Figure 2a, Junction 1), respectively. To further assess isoform-specific genetic regulation, we first computed the Proportion-Spliced-In (PSI) values for each AS event using ASpli (58) using GTEx data, and we identified splicing quantification trait loci (sQTLs) within a 20-kb region surrounding the *CETP* locus (Methods). Notably, SNPs within LD block B-4, which encompassed significant *CETP-203* eQTLs (Figure 1b) exhibit strong associations with AS1 (Figure 2b), primarily in visceral adipocytes, breast, spleen and thyroid tissues, but no tissue-specific effects were detected, since we did not detect significant heterogeneity of its estimate across all tissues (Methods, Supplementary text). The overlap between these sQTLs and *CETP-203* eQTLs indicates that alternative exon 1 quantification, which captures a change in the proportion of alternative exon 1, effectively captures *CETP-203* expression levels. Considering that the expression of CETP-202 may also be captured through the measurement of the alternative splicing event AS9, we detected sQTL for AS9 in multiple tissues (Figure 2b), including adipose tissues, pituitary gland, spleen and thyroid. As for AS1, no tissue-specific effects are observed (Supplementary text). The signals are primarily located at the end of the LD block B-4 (Figure 2b), with the previously reported rs5883 SNP displaying the most significant association (51–53), which is however not identified as an eQTL of CETP.

**Figure 2.**
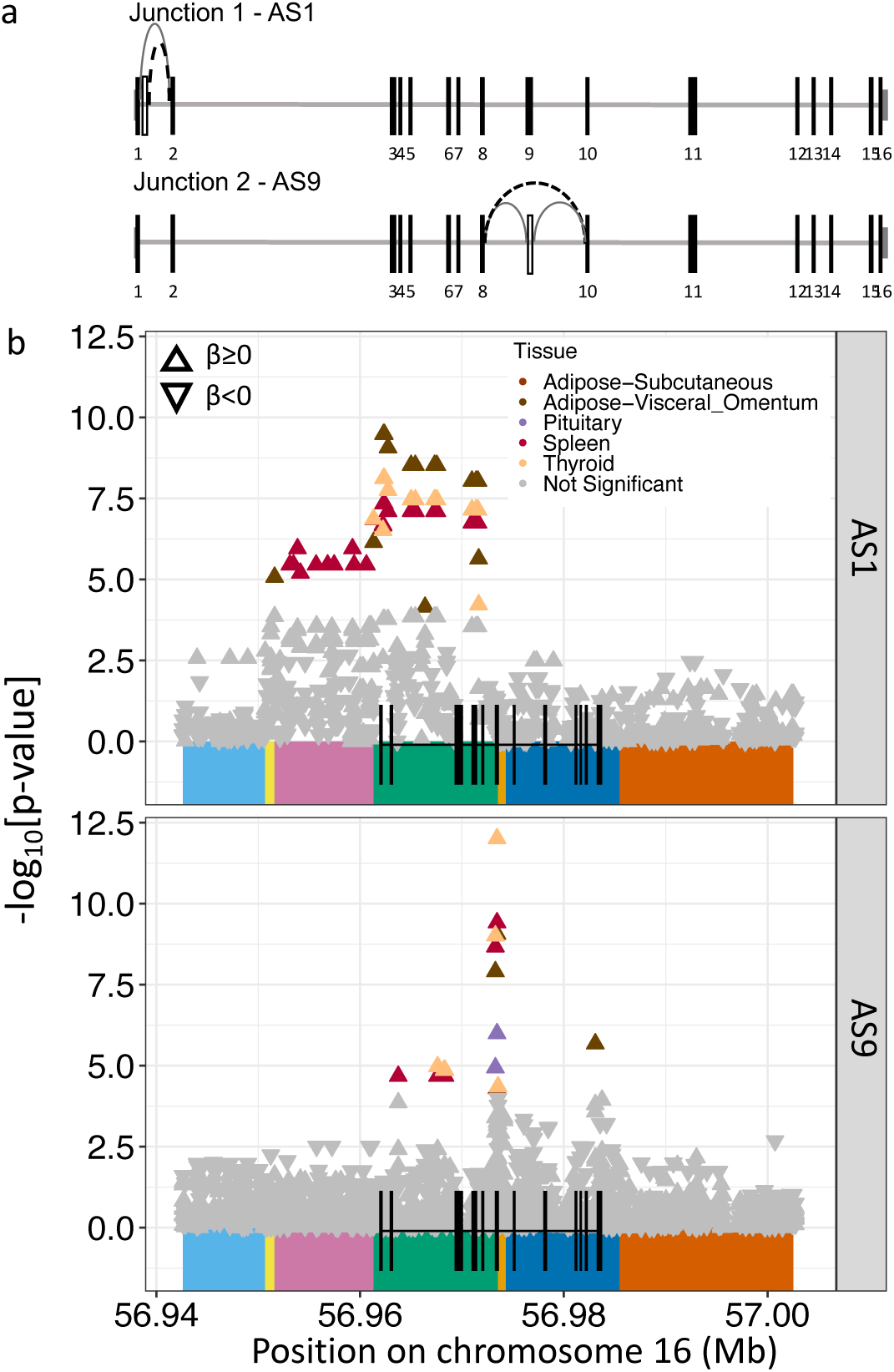
Genetic control of alternative splicing at the *CETP* locus. (a) Diagram of the two splicing junctions in *CETP* gene, where empty boxes represent spliced exons, and the dashed black lines represent the splicing junction variation characterized in ASpli analyses. PSI values represent the proportion of reads associated with the dashed black lines to the total number of reads for both the dashed and full lines. (b) sQTL of the alternative exon 1 (AS1) and alternative splicing of exon 9 (AS9) in the gene *CETP* in different tissues from GTEx data. LD blocks from Figure 1a are represented below the graph. Only tissues with SNPs passing the significance threshold (p-value<0.05/[49 tissues x 7 LD blocks]) are colored. *CETP* gene and its 16 exons are represented at the bottom of each plot.

### Causal Relationships and Tissue-Specific Effects of *CETP* Isoforms on Cardiovascular Disease Phenotypes through Alternative Splicing Analysis

The CETP protein has been extensively implicated in cardiovascular disease (CVD) (21–24), probably due to its impact on the concentration of cholesterol in LDL particles (LDL-C). To investigate the causal relationship between each isoform and CVD-associated phenotypes, we performed Mendelian Randomisation (MR) analyses utilizing eQTLs of gene-level *CETP* expression and sQTLs of AS1 and AS9 as instruments, and coronary artery disease (CAD), as well as HDL-C, LDL-C and TG levels as outcomes (Methods). We employed multivariable models (Methods, Supplementary Figure 5) allowing for isoform-specific contributions while controlling for gene-level expression. When the causal effects of both AS events on a phenotype are consistently in the same direction, *CETP-201* expression variation is likely the influential factor for the phenotype, since it lacks AS1 and AS9. However, if the causal effects of AS1 and AS9 are reversed, either *CETP-202* or *CETP-203* variation is considered the most impactful, as they display only one of the splicing events.

We used the significant eQTLs and sQTLs found in the thyroid tissue, an organ that displays high expression of all three isoforms (Supplementary Figure 1) and strong associations for all three exposures: gene-level *CETP* expression, AS1 and AS9 variation (Figures 1 and 2). *CETP* expression causally modulates all outcomes, as expected (1,2,21–24,40). We also found that an increase in the proportion of both AS1 and AS9 increases the HDL-C concentration, as well as a decrease in LDL-C, TG, and CAD occurrence (Figure 3a), robust to the adjustment for gene-level *CETP* expression. The results remained significant in a sensitivity analysis accounting for directional pleiotropy using the MR-Egger models (Supplementary text, Supplementary Figure 7)(59). Since the effects of both AS events align, our findings indicate that variations in *CETP-201* proportions impact these phenotypes. Specifically, an increase in *CETP-201* leads to increased LDL-C, TG and CAD, but decreased HDL-C, which is consistent with existing literature, as it is the only isoform known to be secreted in plasma (6).

**Figure 3.**
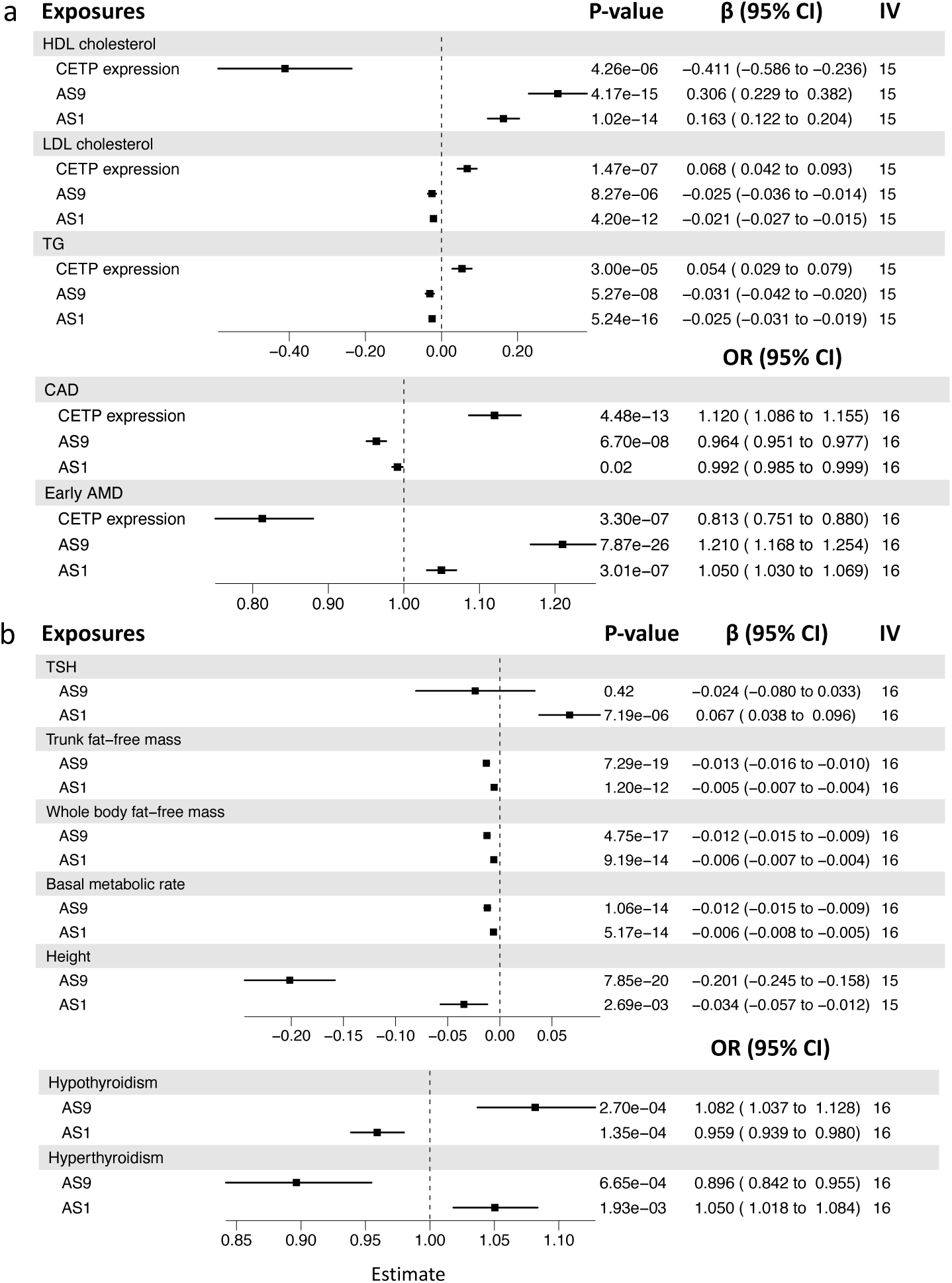
Multivariable Mendelian Randomisation on *CETP* expression and alternative splicing events as exposures and *CETP*-relevant traits as outcomes. Effects of change in the proportion of *CETP* isoforms using multivariable Mendelian Randomisation (MR) on phenotypes (a) previously associated with gene-level CETP expression and (b) associated with thyroid/pituitary glands. Results are from the IVW test, which takes into account gene-level *CETP* expression, alternative exon 1 (AS1) and alternative splicing of exon 9 (AS9). Estimates (Beta) represent the effect of a change of 1 standard deviation on the outcomes. IV represents the number of instrumental variables (IV) used in the analysis. Results for univariable MR are show in Supplementary figures 6 and 9 and for MR-Egger in Supplementary figure 7 and 10.

Given that the transcriptome profile can differ across cell types, potentially affecting isoform dynamics, we sought to investigate whether variation in *CETP* isoform proportion among samples is differentially associated with CAD events across the various tissues available in GTEx (Methods). We report significant associations between AS9 proportion and CAD in adipocyte tissues (P-value_Subcutaneous_=0.01, N=180; P-value_Visceral_=0.04, N=234) and in thyroid (P-value=0.005, N=277) (Supplementary figure 8). The MR results above are consistent with the observed trend in subcutaneous adipocyte tissue, wherein individuals with higher levels of AS9 have a lower proportion of CAD events. Additionally, it is worth reporting that the pattern in liver, which is the main contributor to plasma *CETP*, shows the same trend, although the association is not statistically significant (P-value=0.29) potentially due to a smaller sample size (N=56) and/or cell type heterogeneity in liver bulk RNA-seq. This trend, seen in two tissues that contribute significantly to CETP secretion in plasma (6), could be explained by the fact that an increase of CETP-202 (increase of AS9) decreases the secretion of CETP-201 in plasma (4,41–44). However, we identified an effect in the opposite direction for visceral adipocytes, another tissue involved in plasma CETP secretion, and for the thyroid, where *CETP* expression is the second highest (Supplementary Figure 1) but lacks a known internal function associated with CETP. Altogether, these findings indicate tissue-specific effects of the main isoform on CAD traits, highlighting the complex interplay between isoform-specific regulation and phenotypic outcomes.

### Change in isoform proportion shows causal relationships with phenotypes distinct to *CETP* expression

In addition to CAD, other phenotypes have previously been associated with CETP protein levels or genetic variants within the *CETP* locus in the literature and online databases (Supplementary table 1), such as GWASatlas (60) and PheWeb (61). The list of these phenotypes is reported in Supplementary Tables 1 and 2. For each phenotype, we performed MR to identify the isoform impacting a given phenotype.

*CETP* expression has been found to be associated with early AMD (38,39), where an increase of *CETP* expression is associated with a decrease of the prevalence, which is in the opposite direction to the association with CAD. This opposite effect could thus be due to isoform specific effects. Multivariable MR showed a positive correlation between early AMD and both AS events, robust to controlling for *CETP* expression and pleiotropy (Supplementary text, Supplementary Figures 6 and 7). We observed that variation in AS9 is more significantly associated with early AMD (p-value=7.87 x 10^-26^) than a change in *CETP* expression levels (p-value=3.30 x 10^-7^), and with similar effect sizes in opposite direction, suggesting that variation in the isoform proportion may also be important for this disease.

While the modulation of plasma CETP activity by thyroid hormones and thyroid dysfunction is known(62,63), the specific role of CETP within this tissue, where it is highly expressed, remains unknown. We thus used thyroid dysfunction, specifically results from GWAS on hyperthyroidism and hypothyroidism phenotypes (Supplementary Tables 1 and 2), to evaluate the effect of a change in AS on thyroid function. In multivariable MR models, changes in both AS events were significantly associated with hypothyroidism (p-value_AS9_=2.70 x 10^-4^; p-value_AS1_=1.35 x 10^-4^) and hyperthyroidism (p-value_AS9_=6.65 x 10^-4^; p-value_AS1_=1.93 x 10^-3^) (Figure 3b), but not with *CETP* expression (Supplementary Figure 9). The results for hypothyroidism remained significant after controlling for pleiotropy (Supplementary text, Supplementary Figure 10). Thyroid function is also associated with the thyroid stimulating hormone (TSH), produced by the pituitary gland. We evaluated the causal relationship of AS events on TSH production and found an increase of AS1 to be associated with an increase of TSH levels (p-value=7.19 x 10^-6^) (Figure 3b), robust to controlling for *CETP* expression and pleiotropy (Supplementary Figures 9 and 10). This result suggests that variation in AS1 may impact the production of TSH. Pituitary and thyroid hormones are known to modulate fat-free mass, basal metabolic rate and height(64–66) which are traits that have all been associated with genetic variants in *CETP* in online GWASatlas (60). Based on publicly available GWAS statistics for these phenotypes, we found significant causal relationships only with AS events (Figure 3b) but not with *CETP* expression (Supplementary figures 9 and 10). A change in the proportion of *CETP* isoforms thus have significant effects on anthropometric traits independently of *CETP* expression, potentially through the regulation of pituitary or thyroid hormones. Altogether, these findings indicate that *CETP* isoforms CETP-202 and/or CETP-203 may significantly impact pituitary and/or thyroid glands functions.

### Variation in alternative splicing of exon 9 impacts pulmonary and pregnancy phenotypes

*CETP* has been involved in one of the first sex-specific genetic interactions reported to date in humans, whereby different combinations of genotypes at rs1967309 in *ADCY9* and at rs158477 in *CETP* influenced *CETP* expression in multiple tissues (55). The SNP rs158477, located 188 bp away from the end of exon 9, may impact splicing of this exon, as adjacent regions to spliced exons are known to be involved in splicing regulation (67). To assess whether the epistatic interaction could impact alternative splicing of exon 9, we examined the evidence for interaction between the SNP rs158477 in *CETP* and polymorphisms in *ADCY9* on AS9 variation across tissues in GTEx (Methods).

We found significant interaction effects between rs158477 in *CETP* and several SNPs in *ADCY9* on AS9 (Figure 4a), with the *ADCY9* SNP rs4786452 showing the strongest association (p-value_Interaction_=5.78 x 10^-7^, breast mammary tissue). SNP rs4786452 is in strong LD with rs1967309 (D’>0.99 in the CEU population from 1000 Genomes project (54)) but has a lower minor allele frequency (MAF=0.15 in GTEx, Supplementary text). We note that, because of low MAF at this SNP, one combination of genotypes was missing (CC-rs4786452/GG-rs158477) which can lead to false positive results in interaction analyses. However, the interaction remained significant when merging CC with CG of rs4786452 (p-value_Interaction_=1.70 x 10^-6^) (Figure 4b), which indicates that the presence of a C allele at rs4786452 interacts with genotypes at rs158477 to modulate AS9 variation. These results support the hypothesis that the *ADCY9* locus regulates *CETP* expression and suggest that it may be involved in alternative splicing of exon 9 through an interaction with rs158477.

**Figure 4.**
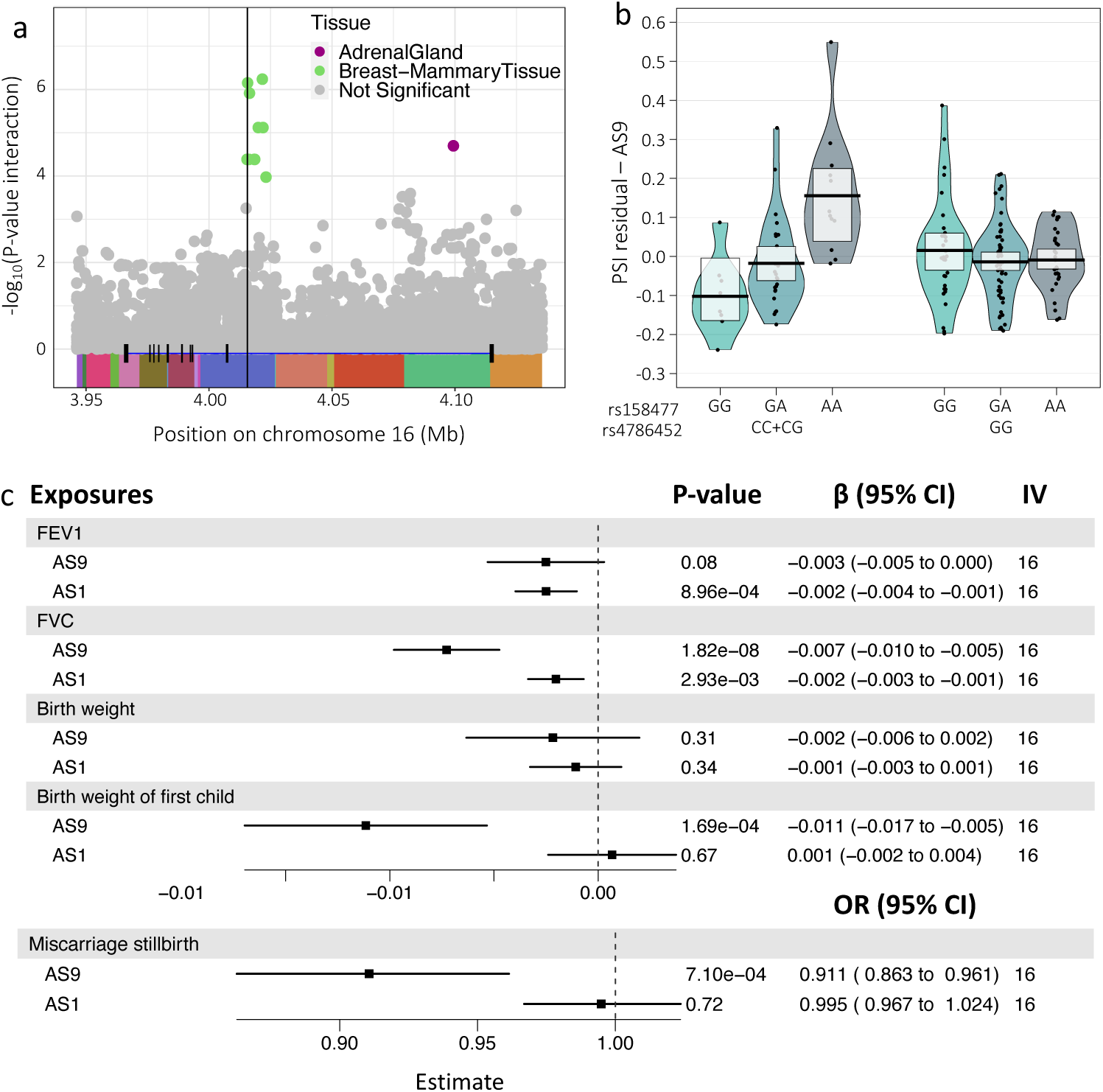
Role of alternative splicing in the epistatic interaction between *ADCY9* and *CETP*. (a) Epistatic interaction between all SNPs in *ADCY9* with MAF>0.05 and the SNP rs158477 in *CETP* on Proportion-Spliced-In (PSI) values for alternative exon 1 (AS1) and alternative splicing of exon 9 (AS9) across tissues in GTEx. Only tissues with SNPs passing significance threshold (p-value<0.05/49) are colored. ADCY9 and its exons are represented at the bottom and colored boxes represent LD blocks estimated with BigLD of gpart package, with a CLQ cut-off at 0.3 for SNPs with MAF>0.05 in the European descent population of GTEx. The vertical black line represents the position of SNP rs1967309. (b) PSI values of AS9 (corrected for covariates used in the regression model) by genotype combination for rs4786452 (*ADCY9*) and rs158477 (*CETP*) in breast mammary tissue. Genotypes CC and CG for rs4786452 were combined for robustness, due to the absence of the CC-rs4786452/GG-rs158477 combination. (c) Multivariable Mendelian Randomization (MR) for AS1 and AS9 exposures on outcomes potentially linked to the previously reported selective pressure in Peruvians. Estimates (Beta) represent the effect of a change of 1 standard deviation on the outcomes. IV represents the number of instrumental variables (IV) used in the analysis. Results for CETP expression and univariable MR are show in Supplementary figure 9 and for MR-Egger in Supplementary figure 10.

The genetic interaction between *CETP* and *ADCY9* was shown to be under selection in the Peruvian population, but the selective pressure remains unknown. However, two pulmonary phenotypes, known to be altered in high-altitude populations (68,69), were found to be significantly impacted by the rs1967309 x rs158477 interaction using the UK biobank data, driven by females (55), namely forced expiratory volume in 1-second (FEV1) and forced vital capacity (FVC). FVC is a measure of lung capacity and indicates restrictive lung disorders, while FEV1 is more commonly used as a measure for obstructive airway disorders. To investigate whether alternative splicing of *CETP* may causally impact these phenotypes, we performed MR analysis on these outcomes (Methods, Supplementary Tables 1 and 2). Our analysis revealed an association between AS9 and FVC (p-value_AS9_=1.82 x 10^-8^), but not with FEV1 (p-value_AS9_=0.08) (Figure 4b, Supplementary figures 9,10).

Another hypothesis put forward to explain the strong selective pattern observed in Peruvians was the impact of CETP modulation in pregnancy or in early life (55). During the early stages of pregnancy, maternal plasma CETP activity and *CETP* expression in the placenta are increased (70–72), and can modulate newborn’s body weight at birth (73). We thus evaluated the impact of AS variation on weight (first child’s weight at birth and the individual’s weight at birth), as well as on stillbirth and miscarriage, through MR (Methods, Supplementary Tables 1 and 2). The weight of the first child was negatively associated with AS9 (p-value=1.69 x 10^-4^) (Figure 4, Supplementary figures 9,10), but not birth weight of the individual (p-value_AS9_=0.31). Furthermore, an increase in the AS9 was associated with a decreased probability of pregnancy complications (miscarriage, stillbirth) (p-value=7.10 x 10^-4^). These results suggest that variation in *CETP* isoform proportions in women is causally associated with pulmonary capacity and pregnancy complications, providing further evidence into the source of the selective pressure acting on *ADCY9* and *CETP*’s combinations of genotypes, potentially modulated through the coregulation of AS9.

## Discussion

In this study, we comprehensively examined the regulation of *CETP* isoforms and identified potential novel functions of CETP. We demonstrated the utility of alternative splicing approaches in estimating isoform levels: although alternative splicing methods typically require high coverage of splicing junctions, resulting in fewer suitable samples for analysis, they offer greater precision in quantification. Furthermore, as each splicing junction only occur in one isoform in *CETP*, combining splicing measurements helped to identify which isoform contributes the most to the effect on a phenotype and allowed us to reveal new associations.

Using multivariable mendelian randomisation, we established significant causal relationships between alternative splicing events of CETP and phenotypes associated with *CETP* expression, such as lipid profile, CAD and early AMD. Notably, since both AS1 and AS9 are consistent in direction, CETP-201 emerged as the most important isoform for these phenotypes in line with its presence in plasma and role in lipid metabolism. These findings support the notion that studies assessing plasmatic CETP or *CETP* expression using gene-level CETP are effectively assessing CETP-201 functions. Furthermore, we showed that changes in the proportion of *CETP* isoforms affected CAD differently across tissues, suggesting tissue-specific and isoform-specific mechanisms of CETP action. Specifically, data from liver and subcutaneous adipose tissue, which are two major producers of plasmatic CETP (6), showed an increase in AS9 that is associated with a decrease in CAD risk, consistent with our MR results. In contrast, the visceral adipose tissue, another major CETP producer, and the thyroid, which has the second highest *CETP* expression (despite not being known as a major contributor of plasma CETP possibly due at least in part to its much smaller size) showed an association in the opposite direction. It raises the question of whether CETP inhibitors, which can enter cells due to their lipophilic nature (32,34), will differentially target each protein isoform. Whether the tissue-specific and isoform-specific effects uncovered here could modulate their potential protective effect against CVD requires further investigation.

AMD is a multifactorial disease caused by damage to the macula. While it involves the accumulation of extracellular lipids, among other factors, the exact pathophysiological changes leading to this accumulation remain to be elucidated (39,74). Cholesterol accumulates in the Subretinal pigmented Epithelial (RPE), which participates in cholesterol exchange with its neighboring tissue, the neural retina, where CETP is expressed (75). Therefore, the process of reverse cholesterol can be crucial to prevent cholesterol accumulation and inflammation. In contrast to the causal relationships observed on CAD, the impact of change in the variation of AS9 on early AMD was similar in effect size to that of *CETP* expression, suggesting significance of isoform proportions in disease development, possibly through lipid homeostasis within the cell (4,41), but future studies should investigate *CETP’s* isoforms in eye tissues. Likewise, our results on CETP isoforms in thyroid and pituitary glands, which are not known to produce plasmatic CETP, suggests an intracellular function for CETP in these tissues, potentially with distinct roles played by different isoforms. Specifically, we found that changes in alternative splicing were causally associated with pituitary and thyroid gland function, as well as many phenotypes influenced by the hormones they secrete.

Previous findings identified a sex-specific epistatic interaction between *ADCY9* and *CETP* genes, specifically with SNP rs158477 located near exon 9 (55). Here, we report an epistatic interaction between *ADCY9* locus and rs158477 on the alternative splicing of exon 9, suggesting that the interaction between both genes may act to modulate CETP isoforms proportions. Interestingly, ADCY9 can activate protein kinase A (PKA), which was found to regulate alternative splicing of other genes (76,77). Furthermore, alternative splicing is known to contribute to adaptive evolutionary changes as it is one mechanism leading to fast response to environmental changes (49). The observed epistatic interaction on AS9 could thus contribute to the coevolution event reported between the two genes. Changes in the proportion of *CETP* isoforms are associated with pregnancy-related phenotypes and pulmonary capacity, both of which are influenced by living at high altitudes (78–82), in line with the selective pressure being observed in the Peruvian population.

There are several limitations in our study. Our analyses investigating the impact of AS changes had limited statistical power given the available number of samples in GTEx, and were restricted to tissues with the highest *CETP* expression, nevertheless generating interesting hypotheses to follow up on. Indeed, replication studies are needed, particularly for the association between change in *CETP* isoforms and CAD occurrence. Additionally, while our analyses focused on identified cis-QTLs, it is important to consider the potential importance of distal regulation, which warrants analyses in larger datasets. This is especially true in the context of the previously identified epistatic interactions involving *CETP*. It is also noteworthy that, in our MR analyses, the three exposures, i.e. *CETP* expression, alternative splicing of exon 9 and alternative exon 1, are not completely independent, which may lead to overcorrection of the effect in the multivariable analysis. To gain a better understanding of the relationship between the three exposures and the phenotypes, it could be beneficial to perform phenotype analysis considering the interaction between the exposures. However, datasets including information of subject with both phenotype and expression data generally have limited statistical power for such analyses, given the limited sample sizes. Furthermore, another limitation is that we were unable to perform sex-specific sQTL analyses due to the limited statistical power resulting from the available number of samples in GTEx. However, the pregnancy-related results from our MR analyses suggest there may be interesting CETP functions specific to women phenotypes. Therefore, conducting follow-up analyses of sex-specific sQTL in larger cohorts is necessary to further explore sex differences in *CETP* regulation.

## Conclusion

In conclusion, our study clearly shows that each *CETP* transcript has its own specific genetic regulation, either directly involving alternative splicing or in their expression modulation through genetic loci. Using these distinct features, we observed that a change in the proportion of its isoforms may have an important impact on phenotypes already known to be associated with *CETP* expression and activity, but also revealed new associations, which suggests a potentially important effect of CETP isoforms in the pituitary and/or thyroid glands. Furthermore, we propose that epistatic interactions involving *CETP* could be mechanistically linked to alternative splicing, which could in turn have importance repercussions on human cardiovascular, pulmonary and reproductive health.

## Acknowledgements

We thank all members of the Hussin lab for their constructive comments and feedback throughout this project. This work was completed thanks to computational resources provided by Compute Canada clusters Graham and Narval. This work was funded by Montreal Heart Institute (MHI) Foundation. Computational work was funded in part by a National Sciences and Engineering Research Council (NSERC) Discovery Grant to JGH (RGPIN-2022-04262). J.G.H. holds a Tier 2 Canada Research Chair (CRC) in responsible multi-omics data science, M.P.D. holds a Tier 1 CRC in precision medicine data analysis. J.C.T. holds a Tier 1 CRC in personalized and translational medicine. I.G. received a PhD scholarship from the MHI Foundation and M.A.L. held a PhD scholarship from Canadian Institutes of Health Research.

## Methods

### CETP transcript definitions

Gene-level CETP (ENSG00000087237) refers to the total gene expression, containing all isoforms count during its quantification. CETP-201 (ENST00000200676.8) contains all 16 exons of CETP. CETP-202 (ENST00000379780.6) has the same exons as CETP-201, except for alternative splicing of the 9th exon. CETP-203 (ENST00000566128.1) differs from CETP-201 only by an alternative exon 1.

### Datasets

We used two datasets for which we had both RNA-seq data and genotyping. The first dataset is obtained from the Genotype-Tissue Expression v8 (GTEx) (83), accessed through dbGaP (phs000424.v8.p2, dbgap project #19088), which includes RNA sequencing across 54 tissues and 948 donors with genetic information available. The cohort contains mainly of European descent (84.6%), ranging in age from 20 to 79 years old. Due to the small sample sizes of GTEx, we applied PCA and ethnicity-based filtering methods to keep the largest homogeneous group, meaning that we only kept self-reported white non-Latino individuals, as described previously (55), resulting in 699 individuals for our analyses, comprising 66 % males and 34 % females. The second dataset used is the GEUVADIS dataset (84) from 1000 Genomes project, which is accessible at https://www.internationalgenome.org/data-portal/data-collection/geuvadis. We kept a total of 287 non-duplicated European samples (CEU, GBR, FIN, TSI).

### BAM processing

For the GTEx (GRCh38) and GEUVADIS (GRCh37) (84) datasets, we extracted the region of *CETP* from the bam files. In GRCh37 (chromosome 16), the region spanned from 56,985,762 to 57,027,757, while in GRCh38 (chromosome 16), it ranged from 56,951,923 to 56,993,845. We performed this extraction using samtools (85), then we used Picard tools (86) to get Fastq files including the unpaired reads. We trimmed the Illumina adaptors from sequencing reads and removed bad quality ends (BQ>20) using TrimGalore! (87). The read files were then mapped to the GRCh38 human genome reference using STAR v2.6.1a (88). During the mapping process, we utilized specific parameters such as outSAMstrandField, outFilterIntronMotifs (RemoveNoncanonicalUnannotated) and outSAMattributes (NH, nM, MD).

### Expression analyses

For the GTEx dataset, pre-processing of expression data was done as described previously (55): briefly, we performed PCA on genetic data, we computed PEER factors (89), transcripts were quantified using RSEM (90) and their expression were normalized using limma (91) and voom (92). Tissues with less than 50 samples were excluded, resulting in samples from 49 different tissues being retained for analysis. In the GTEx dataset, a gene-wide expression quantification quantitative trait locus (eQTL) analysis was performed to assess the association between mutations and gene expression. The genomic region from 56,941,980 to 57,003,666 (20,000bp before and after *CETP* locus) on chromosome 16 (GRCh38) was selected. Positions with MAF below 5% were removed, and only biallelic positions were retained. Indels and SNPs with a Hardy-Weinberg equilibrium p-value below 0.01 were also excluded, resulting in 174 positions for analysis. During the eQTL analyses, two-sided linear regressions on the *CETP* gene expression and its three protein-coding transcripts were done using R (v.3.6.0) (93). Each SNP was coded based on the number of non-reference alleles. The covariates included the first five Principal Components (PCs) computed using FlashPCA2 (94), as well as age, sex, collection site (SMCENTER), sequencing platform (SMGEBTCHT), total ischemic time (TRISCHD), and PEER factors. For the PC analysis, we utilized the imputed genotyping dataset of GTEx v8 using the same filters as mentioned above, and for the filtering based on LD, we followed the recommendations provided by flashPCA2 (https://github.com/gabraham/flashpca). After the filtering process, we retained 100,986 SNPs for performing PCA. The maximum number of PEER factors considered followed the GTEx consortium recommendation based on sample size for each tissue. To assess potential differences in genetic regulation between expression of *CETP* at gene-level and its isoforms, the estimates of their eQTLs were compared using a t-test for each pair (gene-level CETP vs isoform, isoform vs isoform). Only SNPs that passed the threshold of p-value<0.05/[49 tissues * 7 LD blocks] = 0.0001 for at least one of the transcripts in the comparison were considered (Supplementary figure 4).

### Alternative splicing analysis

We estimated Percent Spliced-In (PSI) values using ASpli (58) from the processes BAM files from above. In this study, PSI values estimated by ASpli represent the proportion of reads covering the splicing event, such as alternative exon 1, to the total number of reads covering the junction, including reads associated with alternative exon 1 and regular exon 1 (Figure 2a). The minimum read length was set to 80, and the maximum intron size considered was 10,000 bp. As an alternative approach, we also replicated all results using MAJIQ software (95) (Supplementary text, Supplementary Figures 11 and 12). We note that when using ASpli, the alternative exon 1 was not detected as a splicing event. To address this, we modified the coordinate of the start of the exon 1 of *CETP-203* in the GTF file to ensure overlap with exon 1 of *CETP-201* and *CETP-202*. After estimating PSI values, we filtered out samples with less than 10 reads covering the analyzed splicing junctions. We further restricted the analysis to junctions in tissues with at least 50 samples passing this filter, remaining five tissues for the alternative exon 1 (AS1) and eleven for the alternative splicing exon 9 (AS9) (Supplementary Figure 12). To estimate splicing quantitative trait loci (sQTL), we performed linear regressions using the same positions and covariates as used in the eQTL analyses. These covariates included the first five PCs, age, sex, collection site (SMCENTER), the sequencing platform (SMGEBTCHT), total ischemic time (TRISCHD), and PEER factors. We restricted the number of PEER factors to 15, as was done by GTEx. To assess tissue-specific genetic regulation of alternative splicing, we selected the top sQTL for AS1 and the top sQTL for AS9. We then evaluated if their effects on respective PSI values exhibited heterogeneity across tissue. We considered values to be significant if their p-value, obtained using the metagen function from the meta package in R (96), was below 0.05.

### Epistasis analyses

To examine the interaction between SNPs in *ADCY9* and *CETP* on CETP alternative splicing events, we performed a screening of interaction effects between SNP rs158477 in *CETP* and all biallelic SNPs with MAF>0.05 in the *ADCY9* locus. For the *ADCY9* locus, we kept position from 3,946,204 to 4,135,397 (Chromosome 16, GRCh38, 20 kb around *ADCY9* locus, Supplementary Figure 13). We did the same analysis with SNP rs1967309 in *ADCY9* and all biallelic SNPs with MAF>0.05 in the *CETP* locus (Supplementary text). SNPs with a Hardy-Weinberg equilibrium p-value below 0.01 were excluded. Interactions between SNPs were evaluated using the following model in R: lm(p∼rs1967309*rs158477+Covariates), where the covariates are the same as those used in the splicing QTL analysis without interaction.

### Isoform quantification

Transcript quantifications were estimated using Transcript per million (TPM). The abundance values (TPM) for each CETP isoform were obtained from the GTEx V8 online server. From the remaining 699 individuals, samples with a sum of TPM greater than 0.5 for all three isoforms were selected for the quantification of isoform frequencies (Supplementary Figure 1). The transcript proportions were estimated by combining PSI values from both alternative splicing junctions, meaning AS1 and AS9 (Supplementary Figure 1). The proportions of each transcript were computed using the following formulas:

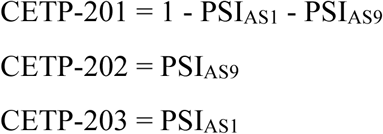

### LD block inference

To infer LD blocks for both *CETP* and *ADCY9* genes, we retained only SNPs in the GTEx dataset with a MAF>0.05 for 20 KB regions surrounding the genes. Using the *BigLD* function from the *gpart* R package (97), we inferred the LD block with a CLQcut of 0.3 and we generated plots for using the *LDblockHeatmap* function from the same package (Figure 1a, Supplementary Figure 13).

### Logistic regression on CAD in GTEx

To evaluate the potential association between alternative splicing and cardiovascular diseases, we conducted a logistic regression analysis in the GTEx dataset. The subjects with CAD were identified based on the variable MHHRTATT (phv00169162.v8.p2) (140 cases) and MHHRTDIS (phv00169163.v8.p2) (126 cases), along with their respective cause of death (Supplementary file 1), using the variable DTHFUCOD (First Underlying Cause Of Death) from GTEx. Individuals were classified as control if they were not a case for both MHHRTATT and MHHRTDIS. In addition, we excluded individuals from the control group if they had the phenotype of “heart disease” (MHHRTDISB (phv00169164.v8.p2)), if their cause of death was associated to the heart but not linked to the two phenotypes mentioned, or if their cause of death was unknown (Supplementary file 2). This resulted in a total of 197 cases and 371 controls included in our analysis. To mitigate potential bias caused by PSI values of 0, we stratified the PSI values in four categories: samples with PSI values of 0 were grouped together, and the remaining samples were stratified into terciles, ensuring similar sample sizes in each group. The logistic regression was performed using this categorial variable instead of the individual PSI values, then compared the model with and without the PSI categories using anova() in R. Since PEER factors may correct for the effect of cardiovascular disease on the transcriptomic profile, we instead calculated 5 SVA (98) on the gene expression protecting the cardiovascular disease status. The other covariables included in the regression analysis were the first five PCs, age, sex, collection site (SMCENTER), the sequencing platform (SMGEBTCHT), total ischemic time (TRISCHD), and we also added *CETP* expression.

### Mendelian Randomization

To assess the causal relationship between a change in the occurrence of alternative splicing events, we performed univariable and multivariable two-sample mendelian randomisation (MR) analyses.

### Exposure Data

For the instrumental variables (IV) of CETP expression, we utilized the results from our gene-level CETP eQTL analysis. For the instruments for alternative exon 1 (AS1) and alternative splicing of exon 9 (AS9), we utilized the results from our sQTL analyses. All participants in the GTEx dataset were of European descent ancestry.

### Outcome Data

We acquired summary statistics from online PheWAS results on European descent individuals, see supplementary tables 1 and 2 for a list of phenotypes and where they were obtained from. There should be no participant overlap between exposure and outcome database. Phenotypes were selected based on literature, observed in online PheWAS results (https://pheweb.sph.umich.edu/, https://atlas.ctglab.nl/), or for other reasons mentioned in results.

### SNP Exclusion Criteria

To determine the IVs for the three exposures, we applied several exclusion criteria. These criteria involved the following thresholds: a derived F-statistic below 10, a p-value above 0.001, and a MAF below 0.01. The derived F-statistic was estimated using an ANOVA model (aov() in R), comparing the model with covariates to the model with covariates and the SNP. We harmonized the exposure and outcome data using the harmonise_data function from the TwoSampleMR package in R (99), and subsequently removed any ambiguous and palindromic SNPs to ensure accurate and reliable results. To assess the correlation structure among the SNPs, we calculated the correlation matrix using the ld_matrix function from the R package ieugwasr (100), considering the 699 individuals of European descent in the GTEx dataset. We pruned high LD (r^2^ > 0.9) pairs of variants using two strategies. In the univariable model, we removed the SNP with the smallest derived F-statistic. In the multivariable model, we avoided removing the SNP with the largest derived F-statistic for each exposure and instead randomly removed one SNP from each pair of SNPs with a correlation exceeding 0.90. Following these filters, only the spleen and thyroid tissues had at least 3 SNPs remaining for each exposure, indicating that these tissues were suitable for further analysis. We performed our analysis in the thyroid tissue since it showed the most significant associations for all three exposures. Detailed summary statistics of the SNPs retained for the thyroid tissue analysis are provided in supplementary tables (Supplementary files 3 and 4).

### Mendelian Randomisation Analysis

We conducted two sample MR analyses using two methods: the Inverse variance weighted (IVW) method and Mendelian randomization Egger regression (MR-Egger) method. There are three core assumptions in MR: i) The variant causes the exposure, ii) there are no confounders of the variant—outcome relationship, iii) the variant does not affect the outcome, except by its effect on the exposure. If the instruments satisfy those assumptions, the IVW methods provides an unbiased estimator of the causal effect. MR-Egger relaxes the conventional instrumental variable assumptions by allowing for directional pleiotropy and is used here as a sensitivity analysis. If the MR-Egger regression intercept coefficient is significantly different from 0 (p-value<0.05), it indicates the presence of directional pleiotropy. In univariable models, the correlation between *CETP* expression, AS1 and AS9 may induce bias due to violations of the 3^rd^ instrumental assumption. To address this, we employed MultiVariable Mendelian Randomization (MVMR) to investigate the causal effect of one exposure on the outcome conditional on the other two exposures. For example, we examined the causal relationship of AS9 by controlling for the effects of CETP expression and AS1. During the MVMR-Egger, we performed this test three times, using each exposure as the reference once to evaluate their intercept. If the intercept was significant in all three tests, it suggested the presence of confounding effects, and the results from MVMR-Egger were considered. If the intercept was not significant for all three tests, we relied on the results from MV-IVW. However, we compared the estimates from MV-IVW with those from MVMR-Egger to ensure consistency. Consistency between IVW and MR-Egger was assessed by examining if the beta coefficients were in the same direction and of comparable magnitude. All MR analyses were conducted using TwoSampleMR and MendelianRandomization (101) R packages. Bonferroni correction was applied to account for multiple testing with a significance threshold of p-value=0.003 (0.05/17 phenotypes).

## References

1. Shinkai H. Cholesteryl ester transfer-protein modulator and inhibitors and their potential for the treatment of cardiovascular diseases. Vasc Health Risk Manag. 2012;8:323–31. doi:10.2147/VHRM.S25238

2. Lagrost L. Regulation of cholesteryl ester transfer protein (CETP) activity: review of in vitro and in vivo studies. Biochim Biophys Acta. 1994 Dec 8;1215(3):209–36. doi:10.1016/0005-2760(94)90047-7 PubMed PMID: 7811705.

3. Haas JT, Staels B. Cholesteryl-ester transfer protein (CETP): A Kupffer cell marker linking hepatic inflammation with atherogenic dyslipidemia? Hepatology. 2015;62(6):1659–61. doi:10.1002/hep.28125

4. Inazu A, Quinet EM, Wang S, Brown ML, Stevenson S, Barr ML, et al. Alternative splicing of the mRNA encoding the human cholesteryl ester transfer protein. Biochemistry. 1992;31(8):2352–8. doi:10.1021/bi00123a021

5. Santana KG, Righetti RF, Breda CN de S, Domínguez-Amorocho OA, Ramalho T, Dantas FEB, et al. Cholesterol-Ester Transfer Protein Alters M1 and M2 Macrophage Polarization and Worsens Experimental Elastase-Induced Pulmonary Emphysema. Front Immunol [Internet]. 2021;12. Available from: https://www.frontiersin.org/articles/10.3389/fimmu.2021.684076

6. Wang Y, van der Tuin S, Tjeerdema N, van Dam AD, Rensen SS, Hendrikx T, et al. Plasma cholesteryl ester transfer protein is predominantly derived from Kupffer cells. Hepatology. 2015;62(6):1710–22. doi:10.1002/hep.27985

7. Jiang XC, Agellon LB, Walsh A, Breslow JL, Tall A. Dietary cholesterol increases transcription of the human cholesteryl ester transfer protein gene in transgenic mice. Dependence on natural flanking sequences. J Clin Invest. 1992;90(4):1290–5. doi:10.1172/JCI115993

8. Oliveira HC, Chouinard RA, Agellon LB, Bruce C, Ma L, Walsh A, et al. Human cholesteryl ester transfer protein gene proximal promoter contains dietary cholesterol positive responsive elements and mediates expression in small intestine and periphery while predominant liver and spleen expression is controlled by 5’-distal sequences. Cis-acting sequences mapped in transgenic mice. J Biol Chem. 1996;271(50):31831–8. doi:10.1074/jbc.271.50.31831

9. Luo Y, Tall AR. Sterol upregulation of human CETP expression in vitro and in transgenic mice by an LXR element. J Clin Invest. 2000;105(4):513–20. doi:10.1172/JCI8573

10. Skoczyńska A, Wojakowska A, Turczyn B, Zatońska K, Wołyniec M, Rogala N, et al. Serum Lipid Transfer Proteins in Hypothyreotic Patients Are Inversely Correlated with Thyroid-Stimulating Hormone (TSH) Levels. Med Sci Monit. 2016;22:4661–9. doi:10.12659/MSM.898134

11. Berti JA, Casquero AC, Patricio PR, Bighetti EJB, Carneiro EM, Boschero AC, et al. Cholesteryl ester transfer protein expression is down-regulated in hyperinsulinemic transgenic mice. J Lipid Res. 2003;44(10):1870–6. doi:10.1194/jlr.M300036-JLR200

12. Hayashibe H, Asayama K, Nakane T, Uchida N, Kawada Y, Nakazawa S. Increased plasma cholesteryl ester transfer activity in obese children. Atherosclerosis. 1997;129(1):53–8. doi:10.1016/s0021-9150(96)06014-5

13. Apostolou F, Gazi IF, Lagos K, Tellis CC, Tselepis AD, Liberopoulos EN, et al. Acute infection with Epstein–Barr virus is associated with atherogenic lipid changes. Atherosclerosis. 2010;212(2):607–13. doi:10.1016/j.atherosclerosis.2010.06.006

14. Lucero D, Zago V, López GI, Graffigna M, López GH, Fainboim H, et al. Does non-alcoholic fatty liver impair alterations of plasma lipoproteins and associated factors in metabolic syndrome? Clin Chim Acta. 2011;412(7–8):587–92. doi:10.1016/j.cca.2010.12.012

15. Izem L, Morton RE. Possible role for intracellular cholesteryl ester transfer protein in adipocyte lipid metabolism and storage. J Biol Chem. 2007;282(30):21856–65. doi:10.1074/jbc.M701075200

16. Zhang Z, Yamashita S, Hirano K, Nakagawa-Toyama Y, Matsuyama A, Nishida M, et al. Expression of cholesteryl ester transfer protein in human atherosclerotic lesions and its implication in reverse cholesterol transport. Atherosclerosis. 2001;159(1):67–75. doi:10.1016/s0021-9150(01)00490-7

17. Radeau T, Lau P, Robb M, McDonnell M, Ailhaud G, McPherson R. Cholesteryl ester transfer protein (CETP) mRNA abundance in human adipose tissue: relationship to cell size and membrane cholesterol content. J Lipid Res. 1995 Dec 1;36(12):2552–61. doi:10.1016/s0022-2275(20)41091-0 PubMed PMID: 8847481.

18. Izem L, Morton RE. Cholesteryl ester transfer protein biosynthesis and cellular cholesterol homeostasis are tightly interconnected. J Biol Chem. 2001;276(28):26534–41. doi:10.1074/jbc.M103624200

19. Greene DJ, Izem L, Morton RE. Defective triglyceride biosynthesis in CETP-deficient SW872 cells. J Lipid Res. 2015;56(9):1669–78. doi:10.1194/jlr.M056481

20. Izem L, Greene DJ, Bialkowska K, Morton RE. Overexpression of full-length cholesteryl ester transfer protein in SW872 cells reduces lipid accumulation. J Lipid Res. 2015;56(3):515–25. doi:10.1194/jlr.m053678

21. Thompson A, Di Angelantonio E, Sarwar N, Erqou S, Saleheen D, Dullaart RPF, et al. Association of Cholesteryl Ester Transfer Protein Genotypes With CETP Mass and Activity, Lipid Levels, and Coronary Risk. JAMA. 2008;299(23):2777–88. doi:10.1001/jama.299.23.2777

22. Webb TR, Erdmann J, Stirrups KE, Stitziel NO, Masca NGD, Jansen H, et al. Systematic Evaluation of Pleiotropy Identifies 6 Further Loci Associated With Coronary Artery Disease. J Am Coll Cardiol. 2017;69(7):823–36. doi:10.1016/j.jacc.2016.11.056

23. Kettunen J, Holmes M V, Allara E, Anufrieva O, Ohukainen P, Oliver-Williams C, et al. Lipoprotein signatures of cholesteryl ester transfer protein and HMG-CoA reductase inhibition. PLoS Biol. 2019;17(12):e3000572. doi:10.1371/journal.pbio.3000572

24. Gautier T, Masson D, Lagrost L. The potential of cholesteryl ester transfer protein as a therapeutic target. Expert Opin Ther Targets. 2016;20(1):47–59. doi:10.1517/14728222.2015.1073713

25. Barter PJ, Caulfield M, Eriksson M, Grundy SM, Kastelein JJP, Komajda M, et al. Effects of torcetrapib in patients at high risk for coronary events. N Engl J Med. 2007;357(21):2109–22. doi:10.1056/NEJMoa0706628

26. Metzinger MP, Saldanha S, Gulati J, Patel K V., El-Ghazali A, Deodhar S, et al. Effect of Anacetrapib on Cholesterol Efflux Capacity: A Substudy of the DEFINE Trial. J Am Heart Assoc. 2020 Dec 15;9(24). doi:10.1161/JAHA.120.018136 PubMed PMID: 33263263.

27. Lincoff AM, Nicholls SJ, Riesmeyer JS, Barter PJ, Brewer HB, Fox KAA, et al. Evacetrapib and cardiovascular outcomes in high-risk vascular disease. New England Journal of Medicine. 2017;376(20):1933–42. doi:10.1056/NEJMoa1609581

28. Schwartz GG, Olsson AG, Abt M, Ballantyne CM, Barter PJ, Brumm J, et al. Effects of dalcetrapib in patients with a recent acute coronary syndrome. N Engl J Med. 2012;367(22):2089–99. doi:10.1056/NEJMoa1206797

29. Nicholls SJ, Ditmarsch M, Kastelein JJ, Rigby SP, Kling D, Curcio DL, et al. Lipid lowering effects of the CETP inhibitor obicetrapib in combination with high-intensity statins: a randomized phase 2 trial. Nature Medicine 2022 28:8. 2022 Aug 11;28(8):1672–8. doi:10.1038/s41591-022-01936-7 PubMed PMID: 35953719.

30. Hartmann G, Kumar S, Johns D, Gheyas F, Gutstein D, Shen X, et al. Disposition into Adipose Tissue Determines Accumulation and Elimination Kinetics of the Cholesteryl Ester Transfer Protein Inhibitor Anacetrapib in Mice. Drug Metab Dispos. 2016;44(3):428–34. doi:10.1124/dmd.115.067736

31. Johns DG, Wang SP, Rosa R, Hubert J, Xu S, Chen Y, et al. Impact of drug distribution into adipose on tissue function: The cholesteryl ester transfer protein (CETP) inhibitor anacetrapib as a test case. Pharmacol Res Perspect. 2019 Dec 1;7(6):e00543. doi:10.1002/prp2.543 PubMed PMID: 31832204.

32. Liu S, Mistry A, Reynolds JM, Lloyd DB, Griffor MC, Perry DA, et al. Crystal Structures of Cholesteryl Ester Transfer Protein in Complex with Inhibitors. J Biol Chem. 2012;287(44):37321–9. doi:10.1074/jbc.M112.380063

33. Rios FJ, Neves KB, Cat AND, Even S, Palacios R, Montezano AC, et al. Cholesteryl Ester-Transfer Protein Inhibitors Stimulate Aldosterone Biosynthesis in Adipocytes through Nox-Dependent Processes. J Pharmacol Exp Ther. 2015 Apr 1;353(1):27–34. doi:10.1124/jpet.114.221002 PubMed PMID: 25617244.

34. Rhainds D, Packard CJ, Brodeur MR, Niesor EJ, Sacks FM, Jukema JW, et al. Role of Adenylate Cyclase 9 in the Pharmacogenomic Response to Dalcetrapib. Circ Genom Precis Med. 2021;14(2):e003219. doi:10.1161/CIRCGEN.121.003219

35. Oliveira HCF, de Faria EC. Cholesteryl ester transfer protein: the controversial relation to atherosclerosis and emerging new biological roles. IUBMB Life. 2011;63(4):248–57. doi:10.1002/iub.448

36. Grion CMC, Cardoso LTQ, Perazolo TF, Garcia AS, Barbosa DS, Morimoto HK, et al. Lipoproteins and CETP levels as risk factors for severe sepsis in hospitalized patients. Eur J Clin Invest. 2010;40(4):330–8. doi:10.1111/j.1365-2362.2010.02269.x

37. Reisinger AC, Schuller M, Sourij H, Stadler JT, Hackl G, Eller P, et al. Impact of Sepsis on High-Density Lipoprotein Metabolism. Front Cell Dev Biol. 2022;9:795460. doi:10.3389/fcell.2021.795460

38. Wang YF, Han Y, Zhang R, Qin L, Wang MX, Ma L. CETP/LPL/LIPC gene polymorphisms and susceptibility to age-related macular degeneration. Sci Rep. 2015;5(1):15711. doi:10.1038/srep15711

39. Cougnard-Grégoire A, Delyfer MN, Korobelnik JF, Rougier MB, Le Goff M, Dartigues JF, et al. Elevated High-Density Lipoprotein Cholesterol and Age-Related Macular Degeneration: The Alienor Study. PLoS One. 2014 Mar 7;9(3):e90973. doi:10.1371/journal.pone.0090973 PubMed PMID: 24608419.

40. Schmidt AF, Hunt NB, Gordillo-Marañón M, Charoen P, Drenos F, Kivimaki M, et al. Cholesteryl ester transfer protein (CETP) as a drug target for cardiovascular disease. Nat Commun. 2021;12(1):5640. doi:10.1038/s41467-021-25703-3

41. Yang TP, Agellon LB, Walsh A, Breslow JL, Tall AR. Alternative Splicing of the Human Cholesteryl Ester Transfer Protein Gene in Transgenic Mice: EXON EXCLUSION MODULATES GENE EXPRESSION IN RESPONSE TO DIETARY OR DEVELOPMENTAL CHANGE (∗). Journal of Biological Chemistry. 1996;271(21):12603–9. doi:10.1074/jbc.271.21.12603

42. Wang S, Deng L, Milne RW, Tall AR. Identification of a sequence within the C-terminal 26 amino acids of cholesteryl ester transfer protein responsible for binding a neutralizing monoclonal antibody and necessary for neutral lipid transfer activity. Journal of Biological Chemistry. 1992;267(25):17487–90. doi:10.1016/S0021-9258(19)37066-8

43. Roy P, MacKenzie R, Hirama T, Jiang XC, Kussie P, Tall A, et al. Structure-function relationships of human cholesteryl ester transfer protein: analysis using monoclonal antibodies. J Lipid Res. 1996 Jan 1;37(1):22–34. doi:10.1016/s0022-2275(20)37632-x PubMed PMID: 8820099.

44. Lira ME, Loomis AK, Paciga SA, Lloyd DB, Thompson JF. Expression of CETP and of splice variants induces the same level of ER stress despite secretion efficiency differences. J Lipid Res. 2008;49(9):1955–62. doi:10.1194/jlr.M800078-JLR200

45. Cunningham F, Allen JE, Allen J, Alvarez-Jarreta J, Amode MR, Armean IM, et al. Ensembl 2022. Nucleic Acids Res. 2022;50(D1):D988–95. doi:10.1093/nar/gkab1049

46. Wright CJ, Smith CWJ, Jiggins CD. Alternative splicing as a source of phenotypic diversity. Nat Rev Genet. 2022;23(11):697–710. doi:10.1038/s41576-022-00514-4

47. Ward AJ, Cooper TA. The Pathobiology of Splicing. J Pathol. 2010;220(2):152–63. doi:10.1002/path.2649

48. Tazi J, Bakkour N, Stamm S. Alternative splicing and disease. Biochimica et Biophysica Acta (BBA) - Molecular Basis of Disease. 2009;1792(1):14–26. doi:10.1016/j.bbadis.2008.09.017

49. Bush SJ, Chen L, Tovar-Corona JM, Urrutia AO. Alternative splicing and the evolution of phenotypic novelty. Philosophical Transactions of the Royal Society B: Biological Sciences. 2017;372(1713):20150474. doi:10.1098/rstb.2015.0474

50. Liu Q, Fang L, Wu C. Alternative Splicing and Isoforms: From Mechanisms to Diseases. Genes (Basel). 2022;13(3):401. doi:10.3390/genes13030401

51. Papp AC, Pinsonneault JK, Wang D, Newman LC, Gong Y, Johnson JA, et al. Cholesteryl ester transfer protein (CETP) polymorphisms affect mRNA splicing, HDL levels, and sex-dependent cardiovascular risk. PLoS One. 2012;7(3):e31930. doi:10.1371/journal.pone.0031930

52. Suhy A, Hartmann K, Newman L, Papp A, Toneff T, Hook V, et al. Genetic variants affecting alternative splicing of human cholesteryl ester transfer protein. Biochem Biophys Res Commun. 2014;443(4):1270–4. doi:10.1016/j.bbrc.2013.12.127

53. Suhy A, Hartmann K, Papp AC, Wang D, Sadee W. Regulation of cholesteryl ester transfer protein expression by upstream polymorphisms: Reduced expression associated with rs247616. Pharmacogenet Genomics. 2015 Jul 22;25(8):394–401. doi:10.1097/FPC.0000000000000151 PubMed PMID: 26061659.

54. Auton A, Abecasis GR, Altshuler DM, Durbin RM, Abecasis GR, Bentley DR, et al. A global reference for human genetic variation. Nature. 2015;526(7571):68–74. doi:10.1038/nature15393

55. Gamache I, Legault MA, Grenier JC, Sanchez R, Rhéaume E, Asgari S, et al. A sex-specific evolutionary interaction between ADCY9 and CETP. Leffler E, Perry GH, Tarazona-Santos E, editors. Elife. 2021;10:e69198. doi:10.7554/eLife.69198

56. Tardif JC, Rhéaume E, Lemieux Perreault LP, Grégoire JC, Feroz Zada Y, Asselin G, et al. Pharmacogenomic determinants of the cardiovascular effects of dalcetrapib. Circ Cardiovasc Genet. 2015;8(2):372–82. doi:10.1161/CIRCGENETICS.114.000663

57. Mizuno A, Okada Y. Biological characterization of expression quantitative trait loci (eQTLs) showing tissue-specific opposite directional effects. European Journal of Human Genetics. 2019;27(11):1745–56. doi:10.1038/s41431-019-0468-4

58. Estefania M, Andres R, Javier I, Marcelo Y, Ariel C. ASpli: Integrative analysis of splicing landscapes through RNA-Seq assays. Bioinformatics. 2021;btab141. doi:10.1093/bioinformatics/btab141

59. Burgess S, Thompson SG. Interpreting findings from Mendelian randomization using the MR-Egger method. Eur J Epidemiol. 2017;32(5):377–89. doi:10.1007/s10654-017-0255-x

60. Watanabe K, Stringer S, Frei O, Umićević Mirkov M, de Leeuw C, Polderman TJC, et al. A global overview of pleiotropy and genetic architecture in complex traits. Nat Genet. 2019;51(9):1339–48. doi:10.1038/s41588-019-0481-0

61. Gagliano Taliun SA, VandeHaar P, Boughton AP, Welch RP, Taliun D, Schmidt EM, et al. Exploring and visualizing large-scale genetic associations by using PheWeb. Nat Genet. 2020;52(6):550–2. doi:10.1038/s41588-020-0622-5

62. Tan KCB, Shiu SWM, Kung AWC. Plasma Cholesteryl Ester Transfer Protein Activity in Hyper-and Hypothyroidism1. J Clin Endocrinol Metab. 1998;83(1):140–3. doi:10.1210/jcem.83.1.4491

63. Duntas LH. Thyroid Disease and Lipids. Thyroid®. 2002;12(4):287–93. doi:10.1089/10507250252949405

64. Shekhda K. The association of hyperthyroidism and immune thrombocytopenia: Are we still missing something? Tzu Chi Med J. 2018 Jul 1;30(3):188–90. doi:10.4103/tcmj.tcmj_139_17

65. Ittermann T, Markus MRP, Bahls M, Felix SB, Steveling A, Nauck M, et al. Low serum TSH levels are associated with low values of fat-free mass and body cell mass in the elderly. Sci Rep. 2021;11:10547. doi:10.1038/s41598-021-90178-7

66. Tarım Ö. Thyroid Hormones and Growth in Health and Disease. J Clin Res Pediatr Endocrinol. 2011;3(2):51–5. doi:10.4274/jcrpe.v3i2.11

67. Barash Y, Calarco JA, Gao W, Pan Q, Wang X, Shai O, et al. Deciphering the splicing code. Nature. 2010;465(7294):53–9. doi:10.1038/nature09000

68. Roh H, Lee D. Respiratory Function of University Students Living at High Altitude. J Phys Ther Sci. 2014;26(9):1489–92. doi:10.1589/jpts.26.1489

69. Weitz CA, Garruto RM, Chin CT. Larger FVC and FEV1 among Tibetans compared to Han born and raised at high altitude. Am J Phys Anthropol. 2016;159(2):244–55. doi:10.1002/ajpa.22873

70. Iglesias A, Montelongo A, Herrera E, Lasunción MA. Changes in cholesteryl ester transfer protein activity during normal gestation and postpartum. Clin Biochem. 1994;27(1):63–8. doi:10.1016/0009-9120(94)90013-2

71. Zhang C, Zhuang Y, Liu X, Chen D, Wang G, Liu Q, et al. Changes in cholesteryl ester transfer protein concentration during normal gestation. European Journal of Lipid Science and Technology. 2006;108(9):730–4. doi:10.1002/ejlt.200600053

72. Gaccioli F, Lager S, Sovio U, Charnock-Jones DS, Smith GCS. The pregnancy outcome prediction (POP) study: Investigating the relationship between serial prenatal ultrasonography, biomarkers, placental phenotype and adverse pregnancy outcomes. Placenta. 2017;59(Suppl 1):S17–25. doi:10.1016/j.placenta.2016.10.011

73. Roland MCP, Godang K, Aukrust P, Henriksen T, Lekva T. Low CETP activity and unique composition of large VLDL and small HDL in women giving birth to small-for-gestational age infants. Sci Rep. 2021;11(1):6213. doi:10.1038/s41598-021-85777-3

74. Fleckenstein M, Keenan TDL, Guymer RH, Chakravarthy U, Schmitz-Valckenberg S, Klaver CC, et al. Age-related macular degeneration. Nat Rev Dis Primers. 2021;7(1):31. doi:10.1038/s41572-021-00265-2

75. Zheng W, Reem RE, Omarova S, Huang S, DiPatre PL, Charvet CD, et al. Spatial Distribution of the Pathways of Cholesterol Homeostasis in Human Retina. PLoS One. 2012;7(5):e37926. doi:10.1371/journal.pone.0037926

76. Gu Q, Jin N, Sheng H, Yin X, Zhu J. Cyclic AMP-Dependent Protein Kinase A Regulates the Alternative Splicing of CaMKIIδ. PLoS One. 2011;6(11):e25745. doi:10.1371/journal.pone.0025745

77. Shi J, Qian W, Yin X, Iqbal K, Grundke-Iqbal I, Gu X, et al. Cyclic AMP-dependent Protein Kinase Regulates the Alternative Splicing of Tau Exon 10. J Biol Chem. 2011;286(16):14639–48. doi:10.1074/jbc.M110.204453

78. Frisancho AR. Human Growth and Pulmonary Function of a High Altitude Peruvian Quechua Population. Hum Biol [Internet]. 1969;41(3):365–79. Available from: https://www.jstor.org/stable/41435777

79. Kiyamu M, Bigham A, Parra E, León-Velarde F, Rivera-Chira M, Brutsaert TD. Developmental and genetic components explain enhanced pulmonary volumes of female peruvian quechua. Am J Phys Anthropol. 2012;148(4):534–42. doi:10.1002/ajpa.22069

80. Nieves-Colón MA, Badillo Rivera KM, Sandoval K, Villanueva Dávalos V, Enriquez Lencinas LE, Mendoza-Revilla J, et al. Clotting factor genes are associated with preeclampsia in high-altitude pregnant women in the Peruvian Andes. The American Journal of Human Genetics. 2022;109(6):1117–39. doi:10.1016/j.ajhg.2022.04.014

81. Pacheco-Romero J, Acosta O, Huerta D, Cabrera S, Vargas M, Mascaro P, et al. Genetic markers for preeclampsia in Peruvian women. Colombia Médica: CM. 2023 Feb 17;52(1):e2014437. doi:10.25100/cm.v52i1.4437

82. Moore LG, Charles SM, Julian CG. Humans at high altitude: Hypoxia and fetal growth. Respir Physiol Neurobiol. 2011;Energetics and Oxygen Transport Mechanisms in Embryos178(1):181–90. doi:10.1016/j.resp.2011.04.017

83. The Genotype-Tissue Expression (GTEx) project. Nat Genet. 2013;45(6):580–5. doi:10.1038/ng.2653

84. Lappalainen T, Sammeth M, Friedländer MR, ‘t Hoen PAC, Monlong J, Rivas MA, et al. Transcriptome and genome sequencing uncovers functional variation in humans. Nature. 2013;501(7468):506–11. doi:10.1038/nature12531

85. Li H, Handsaker B, Wysoker A, Fennell T, Ruan J, Homer N, et al. The sequence alignment/map format and SAMtools. Bioinformatics. 2009;25(16):2078–9. doi:10.1093/bioinformatics/btp352

86. Institute B. Picard Tools [Internet]. 2019. Available from: https://broadinstitute.github.io/picard/

87. Martin M. Cutadapt removes adapter sequences from high-throughput sequencing reads. EMBnet J. 2011;17(1):10–2. doi:10.14806/ej.17.1.200

88. Dobin A, Davis CA, Schlesinger F, Drenkow J, Zaleski C, Jha S, et al. STAR: ultrafast universal RNA-seq aligner. Bioinformatics. 2013;29(1):15–21. doi:10.1093/bioinformatics/bts635

89. Stegle O, Parts L, Piipari M, Winn J, Durbin R. Using probabilistic estimation of expression residuals (PEER) to obtain increased power and interpretability of gene expression analyses. Nat Protoc. 2012;7(3):500–7. doi:10.1038/nprot.2011.457

90. Li B, Dewey CN. RSEM: accurate transcript quantification from RNA-Seq data with or without a reference genome. BMC Bioinformatics. 2011;12(1):323. doi:10.1186/1471-2105-12-323

91. Ritchie ME, Phipson B, Wu D, Hu Y, Law CW, Shi W, et al. limma powers differential expression analyses for RNA-sequencing and microarray studies. Nucleic Acids Res. 2015;43(7):e47–e47. doi:10.1093/nar/gkv007

92. Law CW, Chen Y, Shi W, Smyth GK. voom: precision weights unlock linear model analysis tools for RNA-seq read counts. Genome Biol. 2014;15(2):R29. doi:10.1186/gb-2014-15-2-r29

93. Team RC. R: a language and environment for statistical computing [Internet]. Vienna, Austria: R Foundation for Statistical Computing; 2019. Available from: https://www.R-project.org

94. Abraham G, Inouye M. Fast principal component analysis of large-scale genome-wide data. PLoS One. 2014;9(4):e93766. doi:10.1371/journal.pone.0093766

95. Vaquero-Garcia J, Barrera A, Gazzara MR, González-Vallinas J, Lahens NF, Hogenesch JB, et al. A new view of transcriptome complexity and regulation through the lens of local splicing variations. Valcárcel J, editor. Elife. 2016;5:e11752. doi:10.7554/eLife.11752

96. Schwarzer G, Carpenter JR, Rücker G. Meta-Analysis with R. Use R! [Internet]. Cham: Springer International Publishing; 2015. Available from: https://link.springer.com/10.1007/978-3-319-21416-0 doi:10.1007/978-3-319-21416-0

97. Kim SA, Brossard M, Roshandel D, Paterson AD, Bull SB, Yoo YJ. gpart: human genome partitioning and visualization of high-density SNP data by identifying haplotype blocks. Bioinformatics. 2019;35(21):4419–21. doi:10.1093/bioinformatics/btz308

98. Leek JT, Johnson WE, Parker HS, Jaffe AE, Storey JD. The sva package for removing batch effects and other unwanted variation in high-throughput experiments. Bioinformatics. 2012;28(6):882–3. doi:10.1093/bioinformatics/bts034

99. Hemani G, Zheng J, Elsworth B, Wade KH, Haberland V, Baird D, et al. The MR-Base platform supports systematic causal inference across the human phenome. Loos R, editor. Elife. 2018;7:e34408. doi:10.7554/eLife.34408

100. Elsworth B, Lyon M, Alexander T, Liu Y, Matthews P, Hallett J, et al. The MRC IEU OpenGWAS data infrastructur [Internet]. 2020. doi:10.1101/2020.08.10.244293

101. Yavorska OO, Burgess S. MendelianRandomization: an R package for performing Mendelian randomization analyses using summarized data. Int J Epidemiol. 2017;46(6):1734–9. doi:10.1093/ije/dyx034

