## Supplementary material for "CETP alternative splicing variation impacts human traits": Gamache_et_al.2026.CETP_AS.Supplementary_Text.pdf

### 1. Truncated isoforms in LCL

#### Sashimi plot

Sashimi plots were utilized to visualize splice junctions of the *CETP* gene using the IGV software (1). For the GTEx dataset, all the samples were then merged by tissue, while the European subset were merged in the GEUVADIS dataset, using samtools to merge the samples for plotting in IGV. For the visualization in IGV, we applied a coverage filter of 1000X for a junction in LCL and 2000X for the thyroid gland. This filter allowed us to focus on the junctions associated with the three isoforms of *CETP* (Supplementary Figure 2a).

#### Truncated isoforms

As observed for *CETP* at the gene level, all three isoforms also seem to be expressed across tissues associated to lipid and macrophage with *CETP-201* being the predominant isoform (range of 25.9% to 100%), followed by *CETP-202* (range of 0.0% to 74.1%), then *CETP-203* (range of 0.0% to 35.7%) (Supplementary Figure 1b). Interestingly, cells-EBV-transformed lymphocytes (or Lymphoblastoid Cell Line, LCL) exhibited minimal expression of *CETP-201*. In particular, the cell line showed almost no coverage of exon 1 to exon 6, but it was still possible to detect alternative splicing of exon 9 (Supplementary Figure 2a), representing two unreported *CETP* transcripts. We confirmed that these transcripts were also observed in the LCL from the GEUVADIS dataset (2) This finding may be attributed to a mechanism specific to the transformation process by EBV (3,4). To gain further insights, we investigated the transcription factors associated to EBV infection, obtained from (4) and visualized using the http://epigenomegateway.wustl.edu/ browser (Supplementary Figure 2b). We observed two of its transcription factors, EBNA2 and EBNLP, had enhancer sites in intron 2 and between exons 7 and 10 (Supplementary Figure 2b). During an EBV infection, pathways involved in fatty acid synthesis are induced in newly infected B-cells (5), potentially affecting *CETP* expression, since its activity is increased during infection (6). While many functional domains are retained, the protein structure of these unknown *CETP* transcripts undergoes significant changes, rendering their functions unknown. Further research is needed to investigate these new transcripts, and whether they are found in other tissues, as these isoforms, if present in other tissues, would be masked by the presence of CETP-201 and CETP-202.

### 2. Supplementary results on eQTL and sQTL analyses

#### Residual LD between LD blocks

As we observed significant associations in two LD blocks for gene-level *CETP* and for *CETP-201*, we aimed to determine whether the signals in the less significant LD blocks (B-4) were generated by a residual effect of LD or by a second regulatory region. To investigate this, we included the most significant SNP of B-3 as a covariate in the linear regression analysis of SNPs in B-4. The significance of SNPs within B-4 disappeared, confirming that it is indeed caused by residual LD rather than a second independent regulatory region.

**Tissue-specific regulation of *CETP* expression**

eQTLs exhibiting opposite effects between tissues have been suggested to play a role in the development of complex traits (7). As *CETP* expression has been reported to be tissue-specific (8), confirmed by our results (Supplementary Figure 1), we sought to assess tissue-specificity in the regulation of CETP expression. A single SNP for *CETP-202* barely passed our eQTL threshold in only one tissue (Figure 1b), representing little evidence for CETP-202-specific eQTLs, thus we did not perform tissue-specific analyses for this isoform. We identified the strongest eQTLs for gene-level *CETP, CETP-201* and *CETP-203*: rs56156922 for gene-level *CETP,* rs247616 for *CETP-201* (within LD blocks B-3) and rs711752 for *CETP-203* (within LD blocks B-4). The mean and the standard deviation of the effect size for each eQTL was estimated for each tissue (Supplementary Figure 3). We evaluated the difference of effect sizes for these SNPs for each pair of tissues with an eQTL p-value under 0.0001 in at least one tissue, and then counted the number of tissues for which the SNP showed significant difference (T-test, p-value<0.001 to control for the number of tissue comparisons). For each SNP, we also looked at the overall heterogeneity across all tissues with a Q-cochran test from the package meta (9). We confirmed that eQTL effect sizes differ between tissues (gene-level *CETP*: $\text{p-value}_{\text{Q-Cochran}}<10^{-10}$; *CETP-201*: $\text{p-value}_{\text{Q-Cochran}}<10^{-4}$; *CETP-203*: $\text{p-value}_{\text{Q-Cochran}}<10^{-7}$). *CETP-203* exhibited significant differences in its eQTL effects across tissues at rs711752, however, the effects were consistently in the same direction, suggesting that they may simply indicate stronger statistical evidence in tissues where this isoform is most highly expressed. We describe below the tissue-specific findings at the gene-level and for *CETP-201*.

The effects of rs56156922 on gene-level *CETP* expression in the small intestine is significantly more negative compared to most of the other tissues (Supplementary Figure 3). This observation could be associated with the modulation of *CETP* expression by a fat-rich diet, as the small intestine is responsible for absorbing cholesterol (8). Although not reaching significance as an eQTL in the testis and ovary, we noticed that rs56156922 exhibited opposite directions of effects on CETP expression in these sex-specific tissues compared to other tissues, and these differences were statistically significant (significantly different estimates for 27 and 28 tissue comparisons, respectively). These organs are involved in the production of sexual hormones, and have been involved in differences between sexes for various phenotypes associated with CETP activity, such as HDL-cholesterol levels and cardiovascular disease (10–13). This suggests that the differential regulation of *CETP* in these tissues may play a role in the sex-specific characteristics of these complex traits.

The effects of rs247616 on *CETP-201* expression are significantly different, and in opposite directions, for brain amygdala and whole blood compared to most other tissues (Supplementary

Figure 3). Interestingly, the pattern in amygdala was not detected when looking at gene-level *CETP* expression eQTLs. The amygdala has been associated with Alzheimer's disease (14), and some *CETP* polymorphisms have been associated with this disease, possibly modifying brain structure and neurodegenerative disease susceptibility (15,16). Moreover, amygdala activity has been associated with bone marrow activity, arterial inflammation, and with risk of cardiovascular disease events (17) and atherosclerotic risk (18). These findings suggest a potential association between the *CETP-201* isoform and the development of diseases affecting the amygdala. However, further studies are needed to explore this relationship in more detail.

**Tissue-specificity in alternative splicing**

To assess tissue-specificity regulation of alternative splicing, we identified the strongest splicing quantitative trait loci (sQTL) for alternative exon 1 (AS1) and alternative splicing of exon 9 (AS9). Similar to tissue-specific analyses of eQTLs above, we evaluated the difference of effect sizes for the top sQTL for each splicing event (rs711752 for AS1, rs5883 for AS9) for each pair of tissues with an sQTL p-value under 0.0001 in at least one tissue, and then counted the number of tissues for which the SNP showed significant difference (T-test, p-value<0.001). For each SNP, we also looked at the overall heterogeneity across all tissues with a Q-cochran test from the package meta (9), considered as significant if the p-value were under 0.05. For AS1, the strongest sQTL, rs711752, is also the strongest eQTL for *CETP-203*. This SNP has previously been linked to metabolic syndrome and dyslipidemia (19,20). We did not detect tissue-specific effects for sQTLs in tissues expressing high level of *CETP-203* (Breast-mammary tissue, Spleen, Thyroid, Visceral adipocyte, $\text{p-value}_{\text{Q-Cochran}}$=0.23). Likewise, no tissue-specific effect was observed for rs5883 in AS9 ($\text{p-value}_{\text{Q-Cochran}}$=0.07). This indicates that the genetic regulation of alternative splicing of the exon 9 is likely to be preserved across different tissues.

### 3. Mendelian Randomization Analyses

To better understand the impact of CETP expression at the gene level and the role of isoforms on phenotypes, we employed univariable and multivariable models (Methods, Supplementary Figure 5) using three exposures, namely gene-level *CETP* expression, AS1 and AS9. Univariable models represent the conventional approach of assessing causal relationship between the three exposures and phenotypes independently, while the multivariable model allows to consider the contributions of specific isoforms while controlling for gene-level expression.

Results of multivariable (MV) models are described in the main text. Here, we describe additional results on the associations with gene-level *CETP* expression, as well as comparison of MV models with univariable models and (MV)MR-Egger models for AS1 and AS9 exposures.

**Lipid profile**

Univariable and multivariable IVW models replicated the well-known association between an increase of *CETP* expression with decrease of HDL-c levels, as well as an increase of LDL-c and TG (Supplementary figure 6). AS1 and AS9 are also associated with these outcomes, but with weaker effects and in the opposite direction of *CETP* expression. We performed MR-Egger analysis (Supplementary figure 7) to evaluate the impact of pleiotropy on our results. The intercept indicated significant pleiotropy in the univariable models between AS and HDL-c level. However, after accounting for *CETP* expression in the MV model, the intercept was no longer significant, suggesting that *CETP* expression influences the causal relationship between AS and HDL-c. Furthermore, the estimates between MR-Egger and IVW were in the same direction.

**Diseases known to be associated with CETP expression: CAD and early AMD.**

Causal relationships between *CETP* expression and CAD or early AMD were successfully replicated in univariable and MV analyses (Supplementary figures 6,7). In the univariable models, we observed a nominal significant correlation between increased AS1 and increased risk of early AMD in the IVW test (p-value=0.04) and significant association between increased AS9 and increased risk of early AMD (p-value=7.26 x $10^{-5}$). Those association replicated in the multivariable model (p-value$_{AS1}$=3.01 x $10^{-7}$, p-value$_{AS9}$=7.87 x $10^{-26}$), which included additional instrumental variables, thereby increasing statistical power. Similar to the findings for lipid profiles, the associations between AS1 and AS9 with early AMD were in the opposite direction compared to *CETP* expression.

The intercepts of the MR-Egger analysis were significant for these phenotypes (Supplementary figure 7), and the effect estimates were consistent with those obtained from the IVW method.

**Pituitary and thyroid**

*CETP* was found to be highly expressed in pituitary and thyroid glands. We investigated the impact of changes in *CETP* expression on diseases related to the thyroid (hypo/hyperthyroidism) and the hormone TSH, which is produced by the adrenal gland and can affect thyroid function. Our analysis did not reveal any significant causal relationship between *CETP* expression and these conditions in any of the models (Figure 3, Supplementary figures 9,10). Contrary to multivariable models, univariable models also did not show causal relationship between AS1 or AS9 and thyroid phenotypes after Bonferroni correction, except for a slight association with AS1 for TSH. The intercepts of MR-Egger were not significant.

**Anthropometric traits**

We observed a weakly significant causal relationship between *CETP* expression fat-free mass and basal metabolic rate (BMR) in the univariable models (Supplementary figures 9,10). However, these associations did not persist in multivariable models or MR-Egger analyses. Additionally, body height, which had genome-wide significant associations in the *CETP* locus, did not show any relationship with *CETP* expression. For alternative splicing exposures, in the univariable models,

AS9 showed a negative association with fat-free mass, BMR and body height, which were stronger in the multivariable models, and persisted in MR-Egger (Supplementary figures 9,10). The estimates for AS9 were consistent across all models, indicating robust results. AS1 showed similar effects for fat-free mass and BMR, but not for body height.

**Pulmonary phenotypes**

*CETP* expression showed a strong association with forced expiratory volume in 1 second (FEV1) in both the univariable and multivariable models for IVW and MV-MR-Egger analyses (Supplementary figures 9,10). The univariable MR-Egger model was nominally significant, but the estimate was coherent and the intercept was not significant, indicating no significant horizontal pleiotropy. Associations with forced vital capacity (FVC) were less pronounced and did not persist in the MR-Egger analysis, even in the absence of detected horizontal pleiotropy. This suggests that *CETP* expression, may play an important role in lung function, specifically in relation to obstructive lung diseases, which can be detected by FEV1.

On the other hand, AS9 did not pass the Bonferroni corrected threshold with FVC in the univariable IVW model, but became significant in the multivariable IVW and MR-Egger models (Supplementary figures 9,10), potentially due to increased statistical power. No horizontal pleiotropy was detected by MR-Egger. AS1 association did not pass the Bonferroni corrected threshold for FVC. However, it should be noted that there may also be an association with height, as taller individuals tend to have higher FVC (21).

**Pregnancy-related phenotypes**

On phenotypes related to pregnancy, we found that *CETP* expression is associated with the birth weight of the participant, but in the opposite direction compared to the birth weight of the first child. However, as birth weight is influenced by fetal sex, with male fetuses generally having higher birth weights (22), the opposite direction observed may be due to sex-specific effect. Unfortunately, the available summary data did not allow to stratify by the sex of the participant for birth weight and the sex of the fetuses for the birth weight of the first child.

Alternative splicing (AS1 and AS9) did not show significant associations with the birth weight of the participant: the univariable model showed a suggestive association between AS1 and birth weight, which was lost in the multivariable model. AS9 was not significant in the univariable models (IVW and MR-Egger), but was significant in the multivariable model for birth weight of first child. Once again, the influence of fetal sex could be relevant, but this hypothesis could not be tested with the data at hands.

Regarding the stillbirth/miscarriage phenotypes, neither *CETP* expression nor AS1 showed significant associations. AS9, on the other hand, showed suggestive associations in the univariable models, which became significant in the multivariable IVW model (Supplementary figures 9,10). The intercept of MR-Egger was not significant and the estimates were consistent across the models.

### 4. Epistasis interaction with rs1967309 in ADCY9

Since we observed the effect of a change in alternative splicing of the exon 9 could impact phenotypes, we next evaluated the effect of the interaction between *ADCY9* and *CETP* genes. We detected significant associations between the SNP rs158477 in *CETP* and SNPs within the LD block containing rs1967309 in *ADCY9* on alternative splicing of exon 9 (Figure 4a), with the strongest association with the ADCY9 SNP rs4786452 in breast mammary tissue ($p\text{-value}_{Interaction}=5.78 \times 10^{-7}$), whereas the interaction between rs158477 (*CETP*) and rs1967309 (*ADCY9*) was not significant for neither alternative splicing events in any of the studied tissues, and only nominally significant for interaction with sex in breast mammary tissue with alternative splicing of exon 9 (p-value=0.02). The mutation rs4786452, however, is in high LD with the mutation rs1967309 in almost all populations from 1000 Genomes project (D'>0.99), but with a lower minor allele frequency in breast mammary tissue ($MAF_{rs4786452}=15\%$ vs $MAF_{rs1967309}=38\%$). The C allele of rs4786452 was always on the same haplotype as the A allele of rs1967309, for which we observed an enrichment in the Peruvian population while considering an interaction with rs158477. This suggests an epistasis interaction between rs158477 and ADCY9 locus on AS9.

### 5. Analyses with MAJIQ

MAJIQ is a commonly used software for estimating Percent Spliced-In (PSI) values, but it can introduce complexities in analyzing simple alternative splicing events like AS9. In order to validate the performance of a newer and less well-known software called ASpli, we conducted a comparison of PSI values between the two software tools.

We estimated the PSI values from the processes bam file from above with MAJIQ v2 (23). The configuration files contain the information of the length of RNA-seq reads of 100 and Hg38 reference panel. We did not allow de-novo junctions nor intron retention during the build for MAJIQ. Each tissue had at most three junctions detected. The first one (Junction 1) is for an alternative exon 1, which differentiate isoform *CETP-203* to isoforms *CETP-201/202* and is the same as AS1 from ASpli. The second one (Junction 2) starts at the end of exon 8 and goes to either the beginning of exon 9 or 10. The third (Junction 3) starts at either the end of exon 8 or 9, then finishes at the start of exon 10. The combination of both junctions is similar to AS9 from ASpli.
PSI values were estimated while running MAJIQ with default parameters on all samples separately. 5, 5 and 8 tissues had more than 50 samples with PSI values for junction 1, 2 and 3 respectively, which is less than what was obtained with ASpli. Less samples got a PSI value with MAJIQ, with default parameter, than with ASpli, for which we put a filter to at least 10 reads for a junction instead of the default 5 reads (Supplementary Figure 12).

### PSI values comparison

ASpli and MAJIQ have two distinct approaches to quantified PSI values. While comparing PSI values for both methods, we observed strong correlation for alternative exon 1 (AS1) in ASpli with junction 1 in MAJIQ, and skipping exon 9 (AS9) in ASpli with junction 2 and 3 in MAJIQ (Supplementary Figure 11). Only LCL showed lesser correlation for junction 2 and exon skipping 9, potentially due to the truncated transcript in this cell line (Supplementary Figure 2).
We performed sQTL as we did for ASpli in the main method. Hit regions for sQTL were highly similar between both methods. However, observing the strength of the association with a derived F-statistic, in some tissues, we observed an increase in the strength of the association for AS1 with ASpli, but in other tissues, such as spleen, we observed a slight decrease in the strength of the association for AS9.

In MAJIQ for the junction 1, which was estimated with adipocyte, thyroid, breast and spleen tissues, we did not observed an heterogeneity of the effect of the most significant $sQTL_{AS1}$ rs711752 for this junction ($p\text{-value}_{Q\text{-Cochran}}=0.17$). For junction 2, which was estimated with the same tissues as junction 1, we also did not observe an heterogeneity of the effect of the most significant $sQTL_{AS9}$ rs5883 for this junction ($p\text{-value}_{Q\text{-Cochran}}=0.27$). However, for junction 3, which was estimated with the same tissues than the other two junctions, plus pituitary, lung and LCL, we observed significant heterogeneity of the effect of rs5883 on PSI values ($p\text{-value}_{Q\text{-Cochran}}=0.005$), which was mostly caused by the LCL ($p\text{-value}_{Q\text{-Cochran without LCL}}=0.08$). We did not observe heterogeneity with ASpli caused by LCL, potentially since AS9 is approximately the average from junctions 2 and 3, which could increase the variance of the effect of rs5883 on this junction.

### Supplementary figures

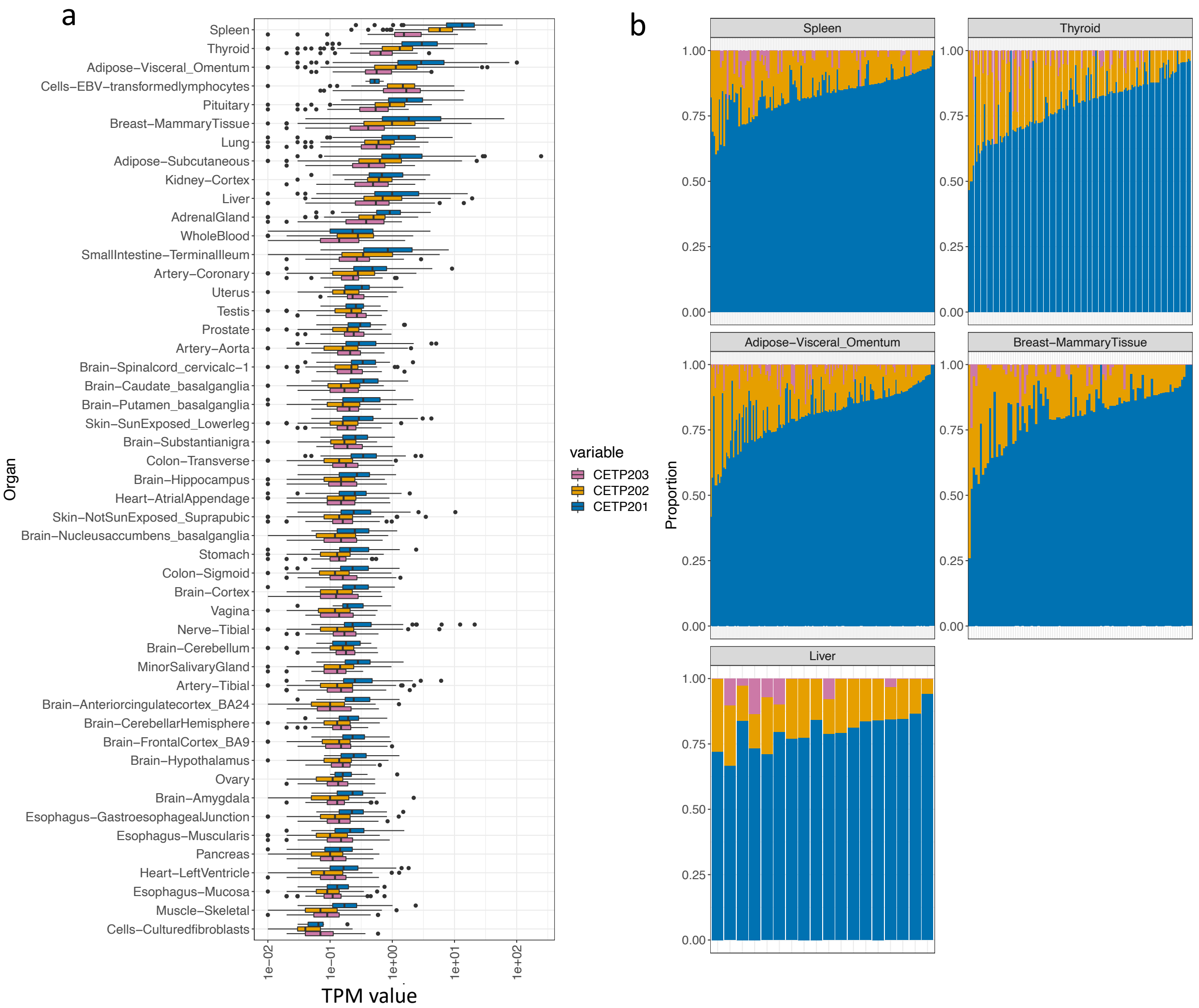


**Supplementary Figure 1. Expression of CETP transcripts by tissue in GTEx dataset.** (a) Transcript per million (TPM) for the three protein coding CETP isoforms generated by RSEM in GTEx. Values of 0 were removed from this graph and x axis was log-transformed. (b) Proportion of each transcript for each sample for 5 tissues, reported using Proportion-Spliced-In (PSI) values estimated by ASpli. Samples are included only if they had non-zero values for alternative exon 1 and alternative splicing of exon 9.

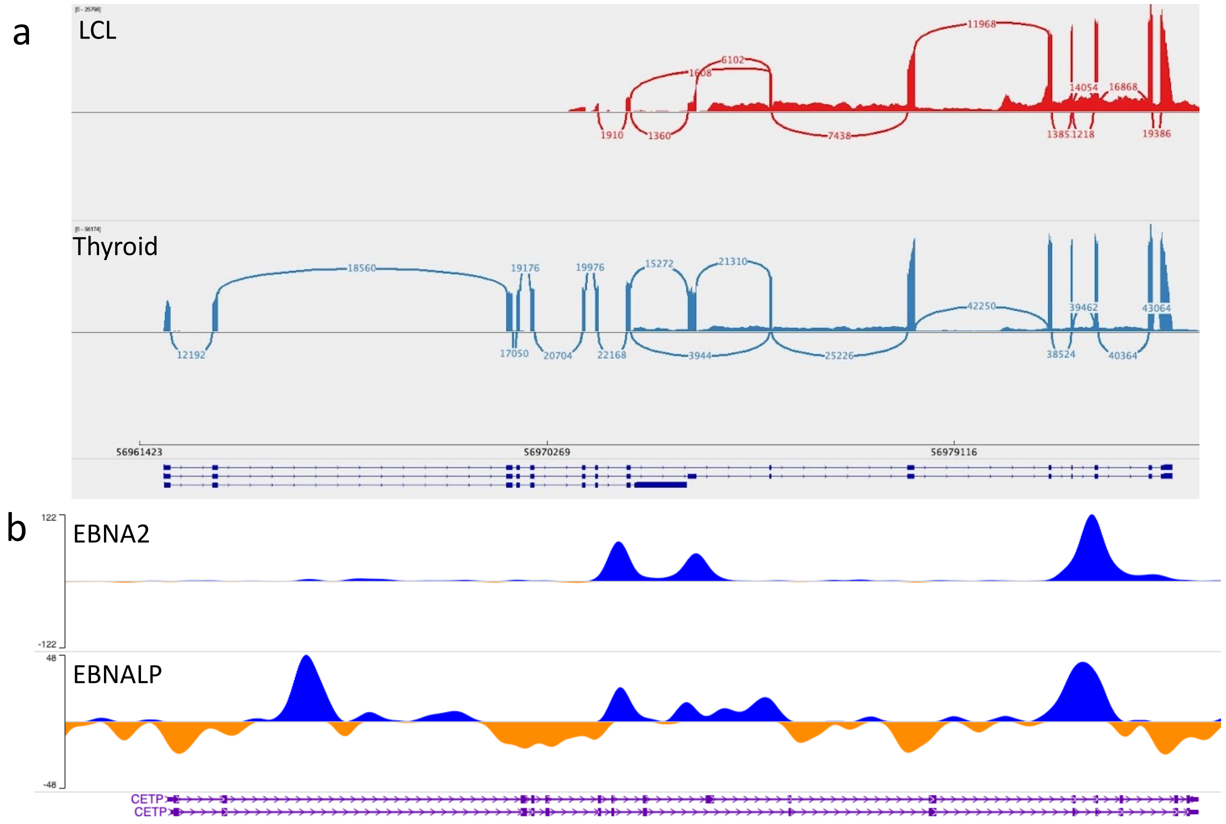


**Supplementary Figure 2. CETP isoform in cells-EBV-transformed lymphocytes (LCL).**
(a) Sashimi plot of the *CETP* gene in the CETP isoform in LCL and thyroid tissue visualized in IGV, made from merging all samples from the LCL (n=174) and thyroid (n=653). Junctions are shown when there was a minimum of 1000 reads junctions for LCL and 2000 for Thyroid. (b) Promoter region of Epstein-Barr virus (EBV) genes from http://epigenomegateway.wustl.edu/browser/ database compared to the coverage of CETP in LCL from GEUVADIS.

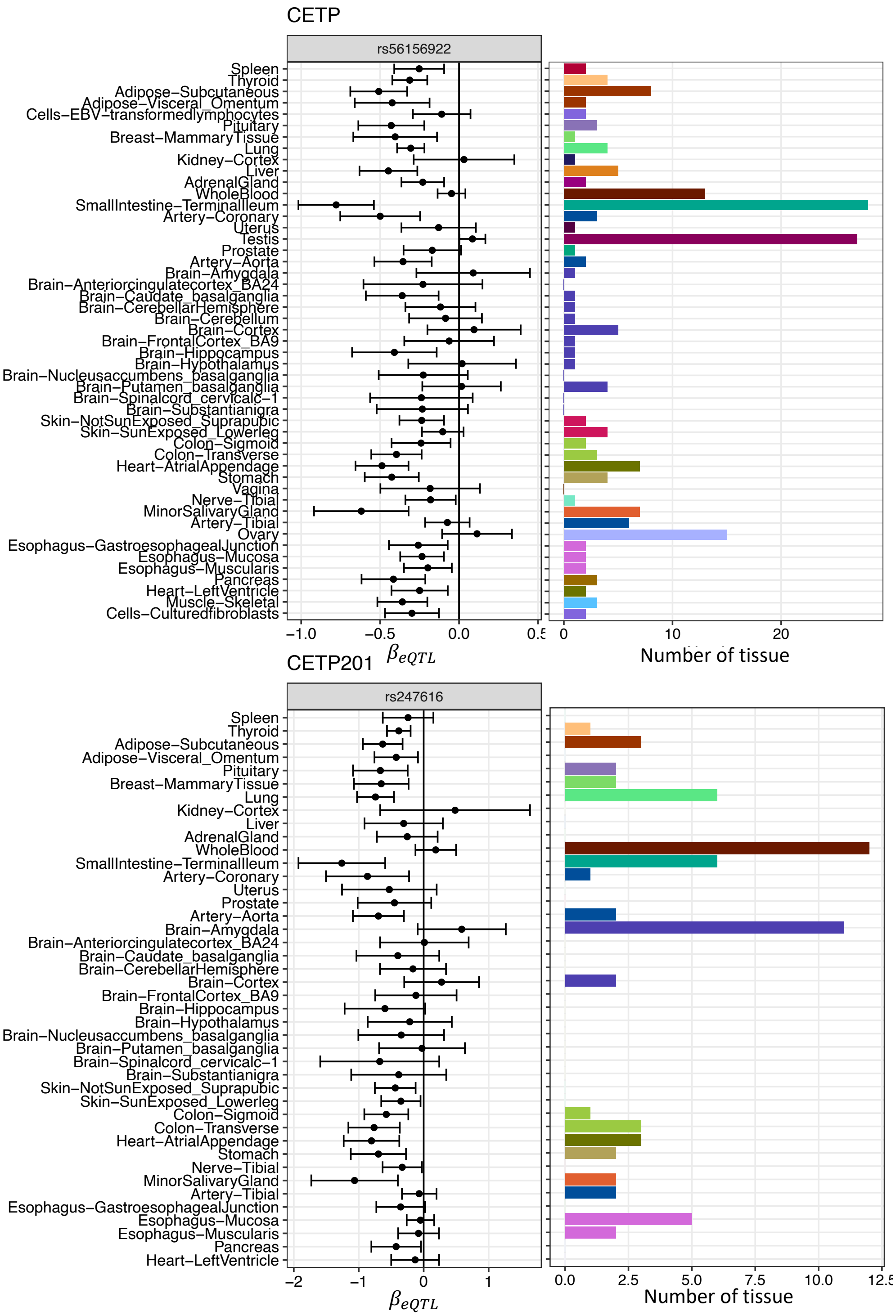


**Supplementary Figure 3. Tissue-specificity of eQTLs for gene-level CETP (top) and CETP-201 (bottom) across tissues.**

The SNPs chosen are the strongest eQTL in the LD block B-3. The barplot represent the number of tissues the effect size estimates significantly differ from the effect size estimate from the labeled tissue. Tissues are sorted according to the gene-level expression of CETP.

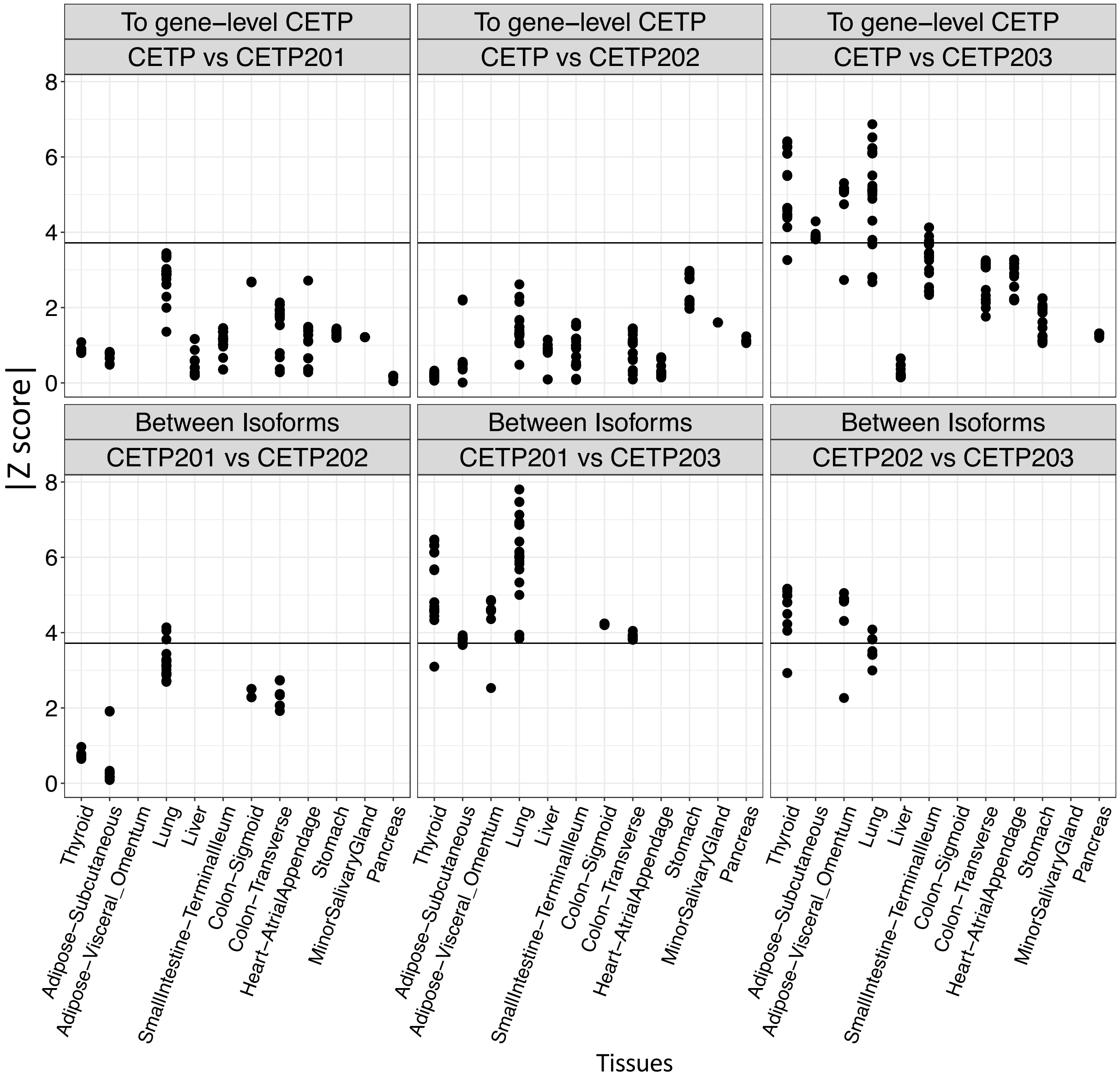


**Supplementary figure 4. Comparison of the effect size estimate of isoform-level with gene-level *CETP* expression (top) and between isoforms (bottom).**

Absolute Z-score of a T-test comparing the effects of all SNPs between two analyses with a p-value < 0.0001 (CETP-wide significance) for at least one analysis in the comparison. Tissues are sorted according to the gene-level expression of CETP. Black horizontal lines represent the Z-score corresponding to a p-value of 0.001.

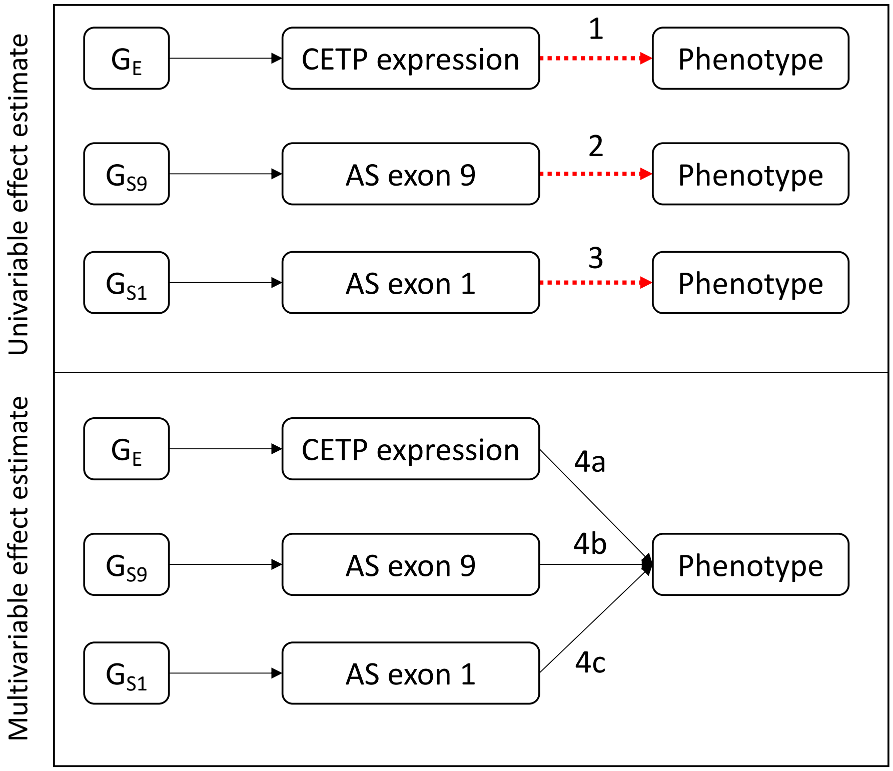


**Supplementary Figure 5. Representation of the effect estimated in the univariable (Top) and multivariable (Bottom) mendelian randomisation (MR) analyses.**

Arrows indicate causal effect studied in each test. Effect estimate 1 is an univariable MR on CETP expression. Effect estimate 2 is an univariable MR on alternative splicing of exon 9. Effect estimate 3 is an univariable MR for alternative exon 1. Effect estimates 4 is a multivariable MR model for which we considered CETP expression, alternative splicing of exon 9 and alternative exon 1. Effect estimate 4a is associated to CETP expression, effect 4b with CETP-202 proportion, 4c to CETP-203 proportion in this model.

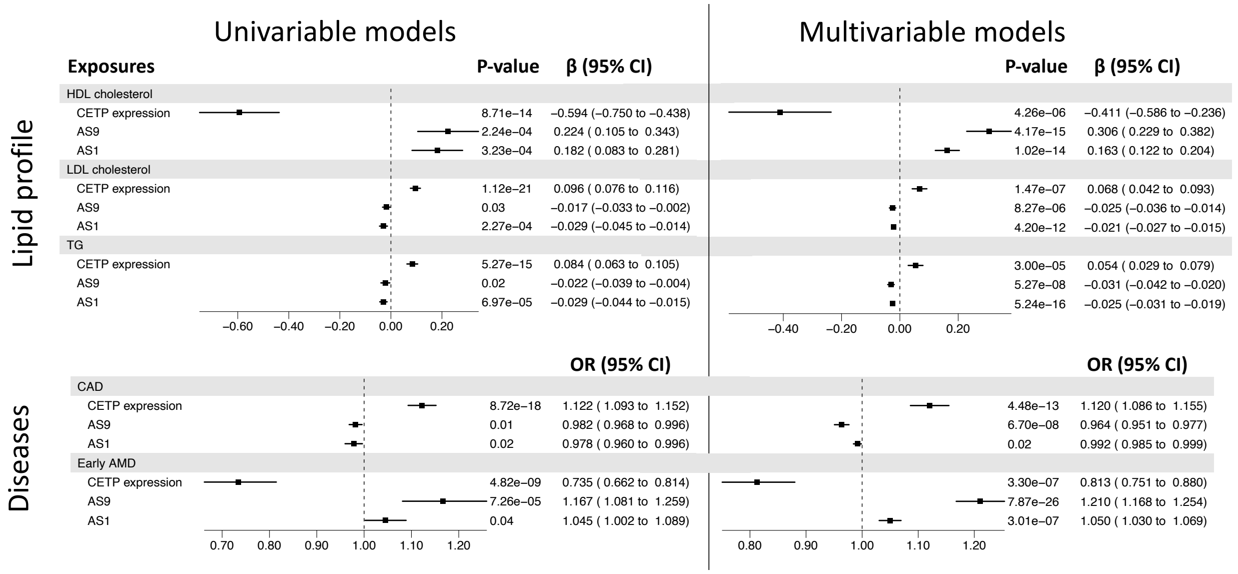


**Supplementary Figure 6. Effects of change in the proportion of CETP isoforms using IVW univariable and multivariable Mendelian Randomisation on phenotypes previously associated with gene-level CETP expression**.

Results are from the IVW test. The multivariable MR takes into account gene-level CETP expression, alternative exon 1 (AS1) and alternative splicing of exon 9 (AS9). Estimates (Beta or Odd Ratio (OR) depending on the phenotype) represent the effect of a change of 1 standard deviation on the outcomes.

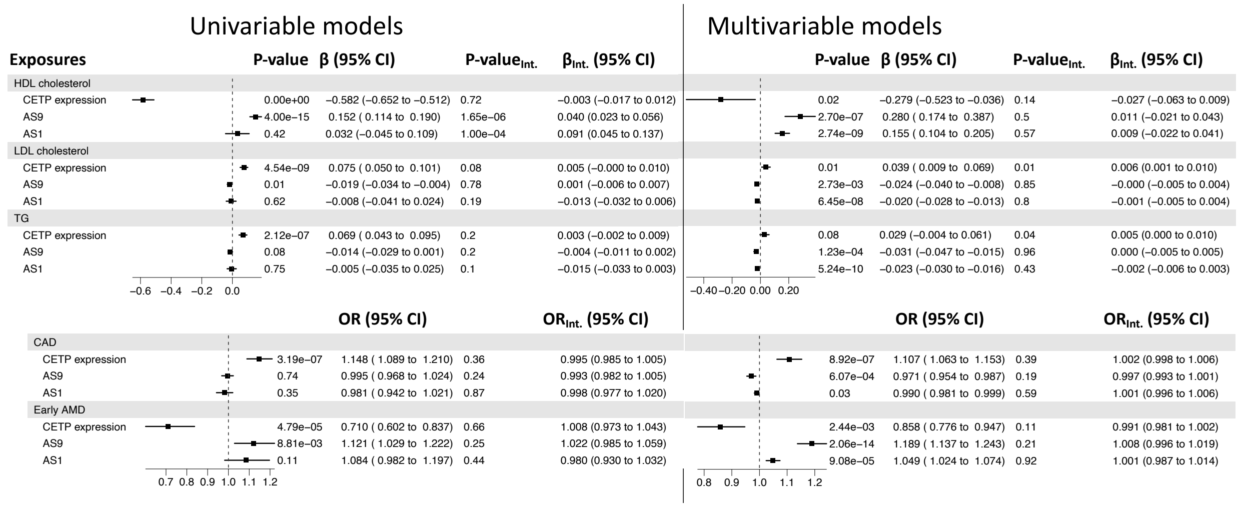


**Supplementary Figure 7. Effects of change in the proportion of CETP isoforms using MR-Egger univariable and multivariable Mendelian Randomisation on phenotypes previously associated with gene-level CETP expression.**

Results are from the MR-Egger test. The multivariable MR takes into account gene-level CETP expression, alternative exon 1 (AS1) and alternative splicing of exon 9 (AS9). Estimates (Beta or Odd Ratio (OR) depending on the phenotype) represent the effect of a change of 1 standard deviation on the outcomes. P-value and estimate (Beta or OR) of the intercept (Int.) obtain by MR-Egger indicate the presence or absence of horizontal pleiotropy in the model.

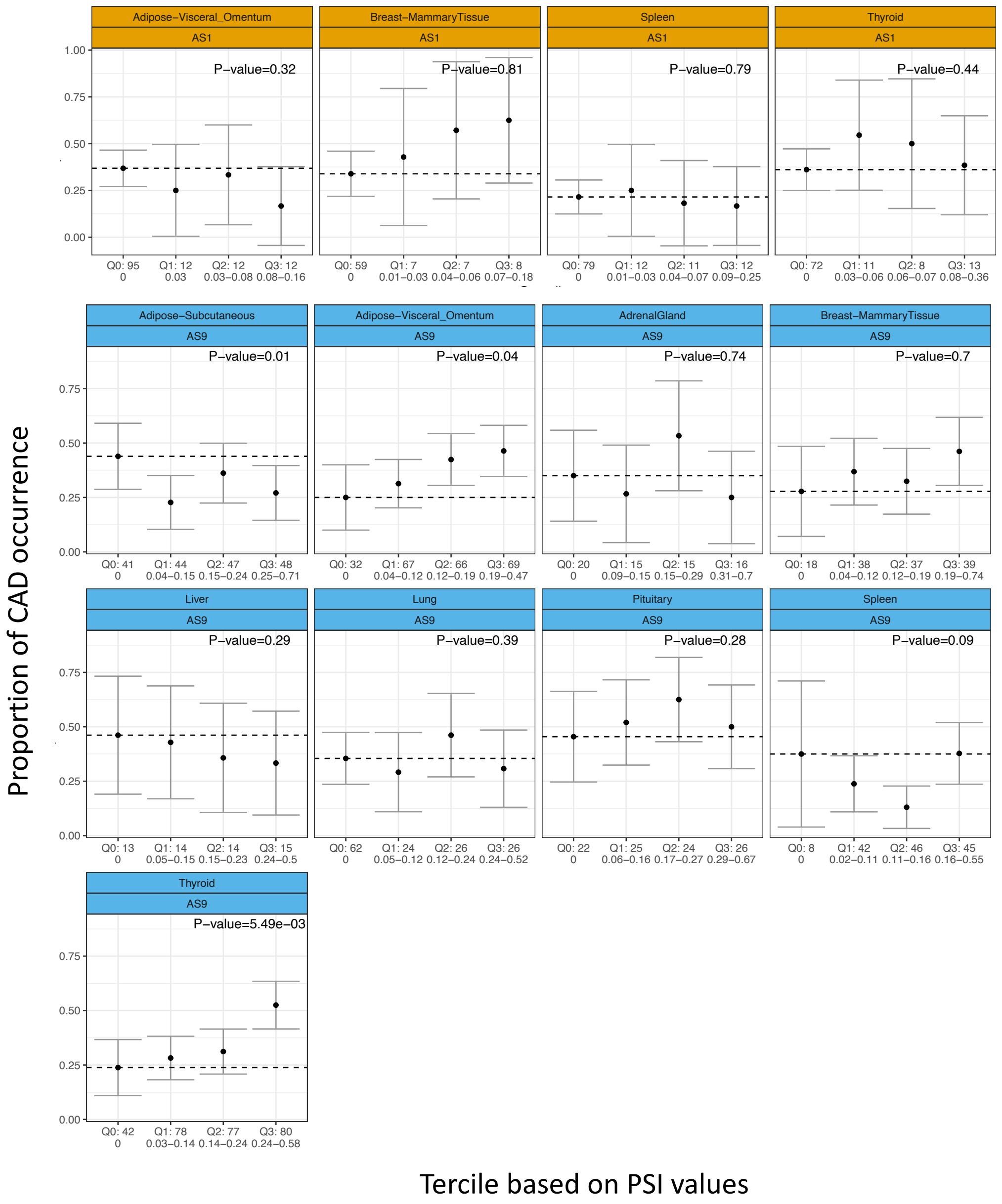


**Supplementary figure 8. Relationship between Proportion-Spliced-In (PSI) values and proportion of coronary artery disease (CAD) occurrence in GTEx individuals.**

PSI values were separated in four group (Q0 : PSI values equal 0; Q1 is the first tercile, Q2 the second and Q3 the third). The proportion of individuals with cardiovascular events in each group of PSI values for alternative exon 1 (AS1) alternative splicing of exon 9 (AS9), shown by tissues. Black dashed lines represent the proportion of Q0. P-value reported for each tissue results from a logistic regression model on CAD, using tercile group as categorial variable. The numbers on the x axis represents the number of samples in this group and the interval is the range of PSI values in the group.

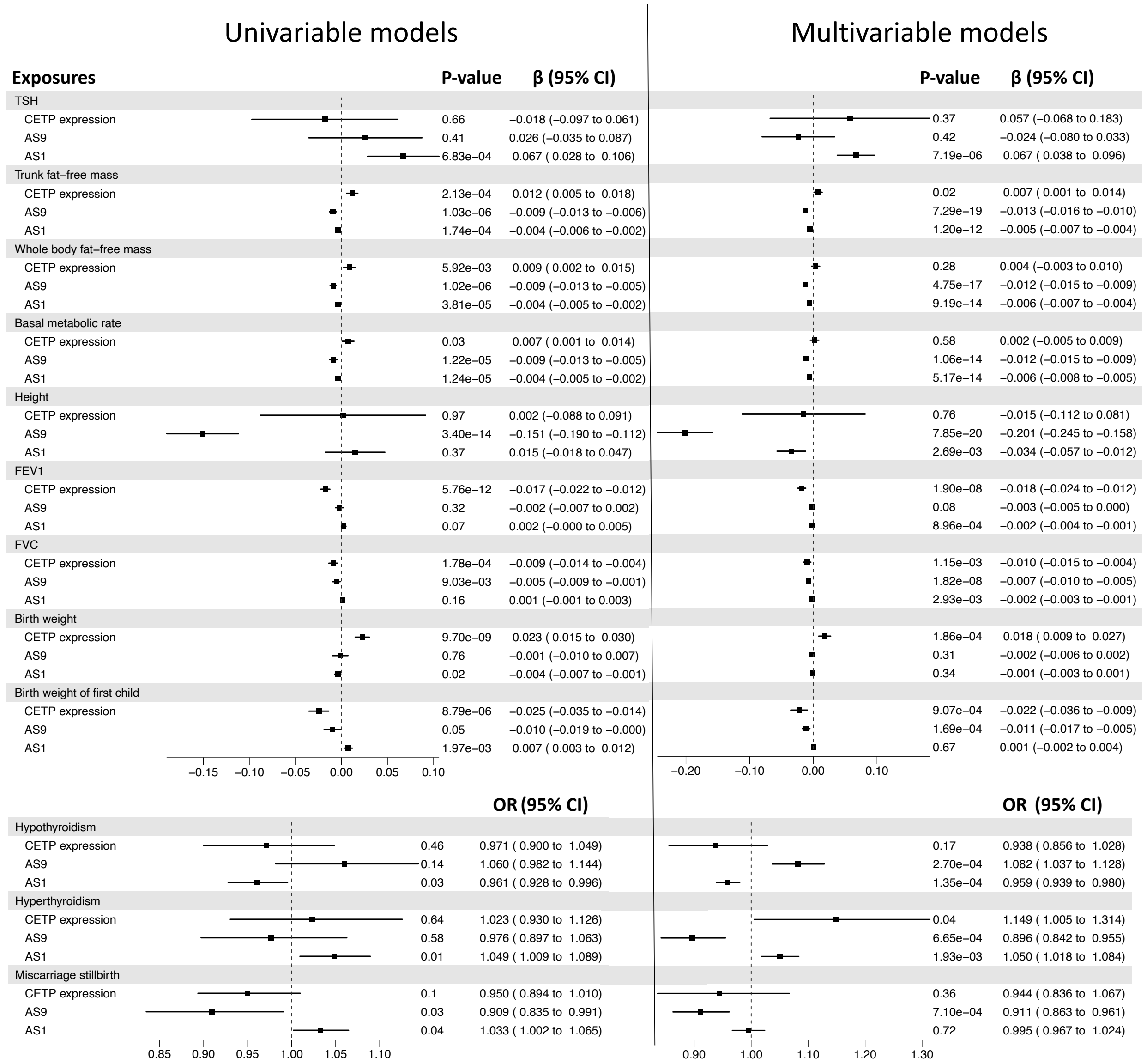


**Supplementary Figure 9. Effects of change in the proportion of CETP isoforms using IVW univariable and multivariable Mendelian Randomisation on phenotypes associated with thyroid/pituitary gland or potentially under selective pressure.**

Results are from the IVW test. The multivariable MR takes into account gene-level CETP expression, alternative exon 1 (AS1) and alternative splicing of exon 9 (AS9). Estimates (Beta or Odd Ratio (OR) depending on the phenotype) represent the effect of a change of 1 standard deviation on the outcomes.

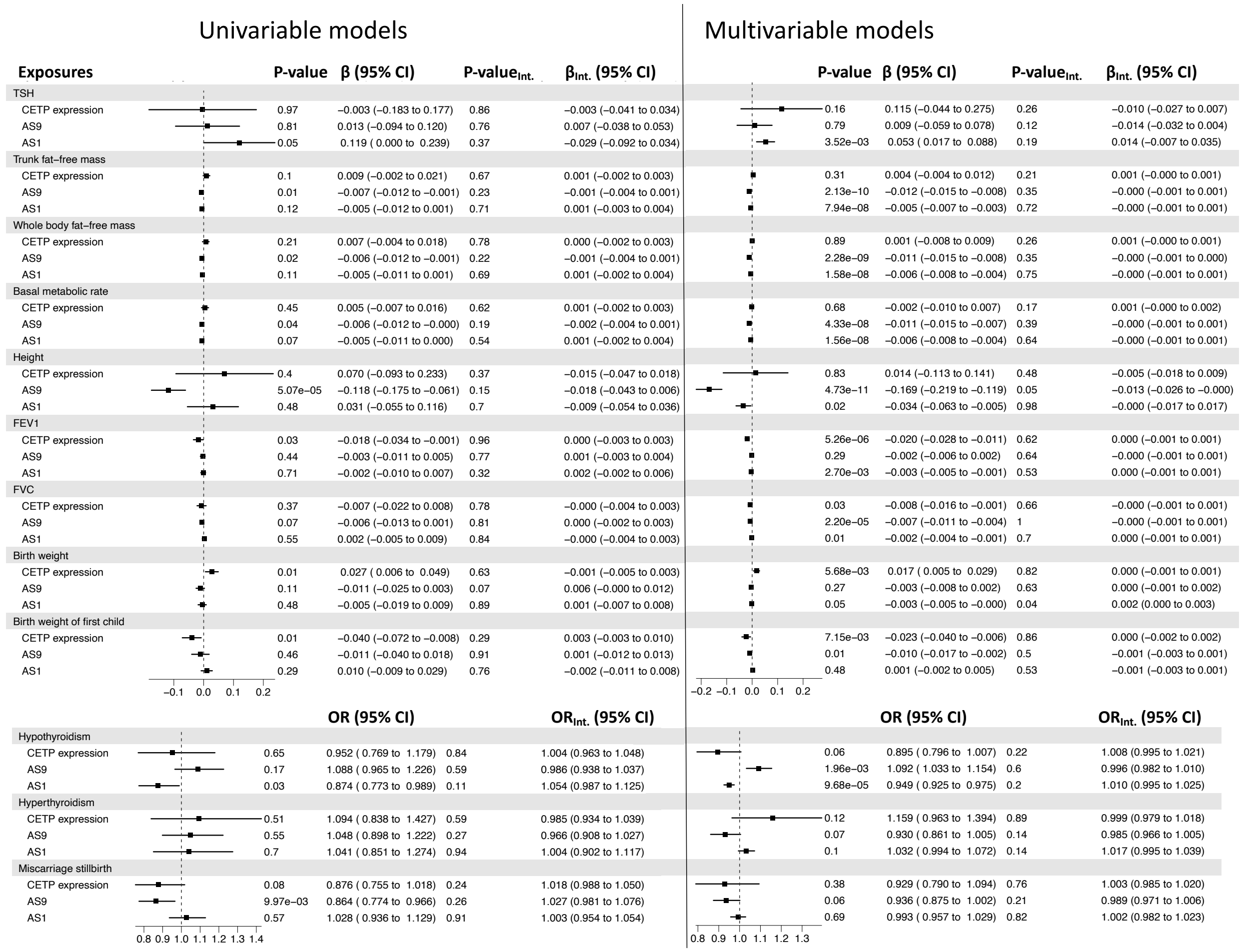


**Supplementary Figure 10. Effects of change in the proportion of CETP isoforms using MR-Egger univariable and multivariable Mendelian Randomisation on phenotypes associated with thyroid/pituitary gland or potentially under selective pressure.**

Results are from the MR-Egger test. The multivariable MR takes into account gene-level CETP expression, alternative exon 1 (AS1) and alternative splicing of exon 9 (AS9). Estimates (Beta or Odd Ratio (OR) depending on the phenotype) represent the effect of a change of 1 standard deviation on the outcomes. P-value and estimate (Beta or OR) of the intercept (Int.) obtain by MR-Egger indicate the presence or absence of horizontal pleiotropy in the model.

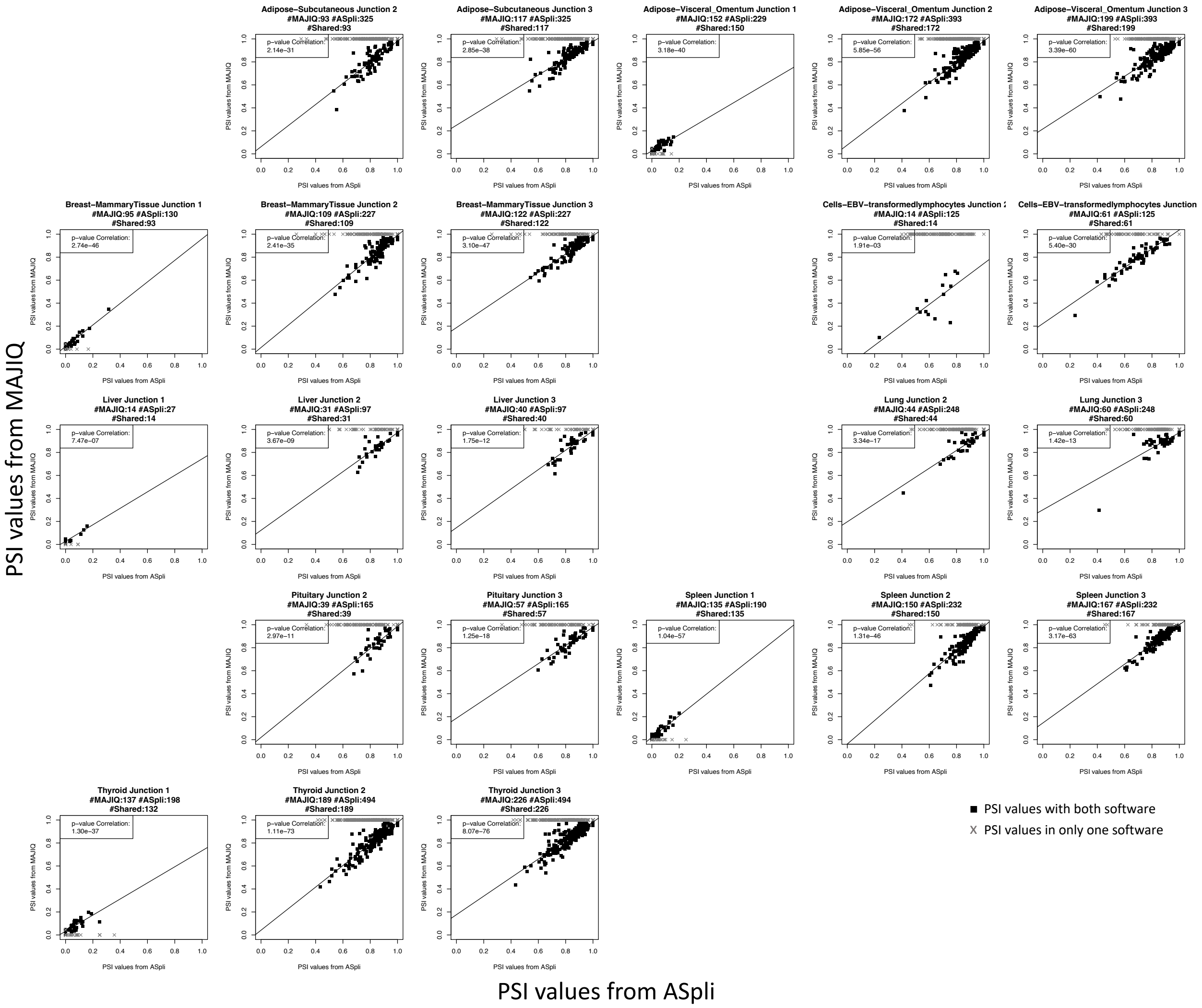


**Supplementary Figure 11. Comparison of Percent Spliced-In (PSI) values obtain by MAJIQ (y axis) and ASpli (x axis) softwares.**

Junction 1 represents the splicing junction at the beginning of exon 2, quantifying alternative of exon 1 (AS1). Junction 2 represents the splicing junction at the end of exon 8, and junction 3 represents the splicing junction at the beginning of exon 10, both quantifying alternative splicing of exon 9 (AS9). Gray values indicate samples without PSI values in MAJIQ using default parameters, but with values in ASpli.

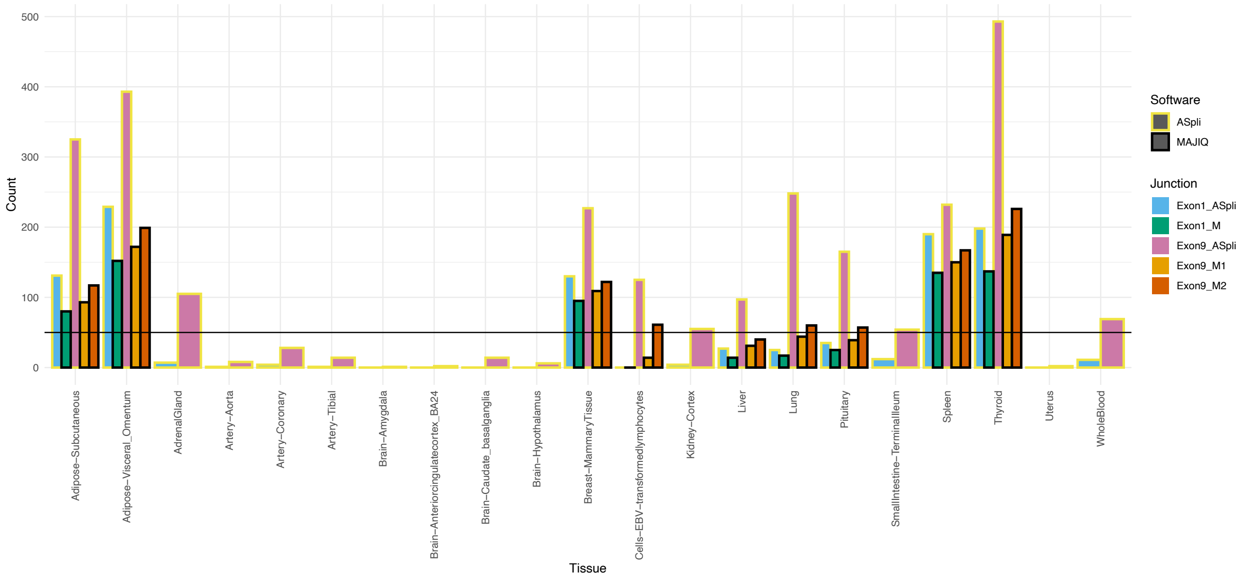


**Supplementary Figure 12. Number of samples with PSI values obtained from MAJIQ and ASpli.**

MAJIQ was used with default parameters, while samples with less than 10 reads of coverages for the junction were filtered out for ASpli. Alternative of the exon 1 are named Exon1_”Sofware” and alternative splicing of the exon 9 are named Exon9_”Software”, where “Software” is either MAJIQ (M : AS1, M1 : Junction represents the splicing junction at the end of exon 8, M2 : Junction at the beginning of exon 10) or ASpli. Horizontal black lines indicate the threshold of 50 samples per junction used.

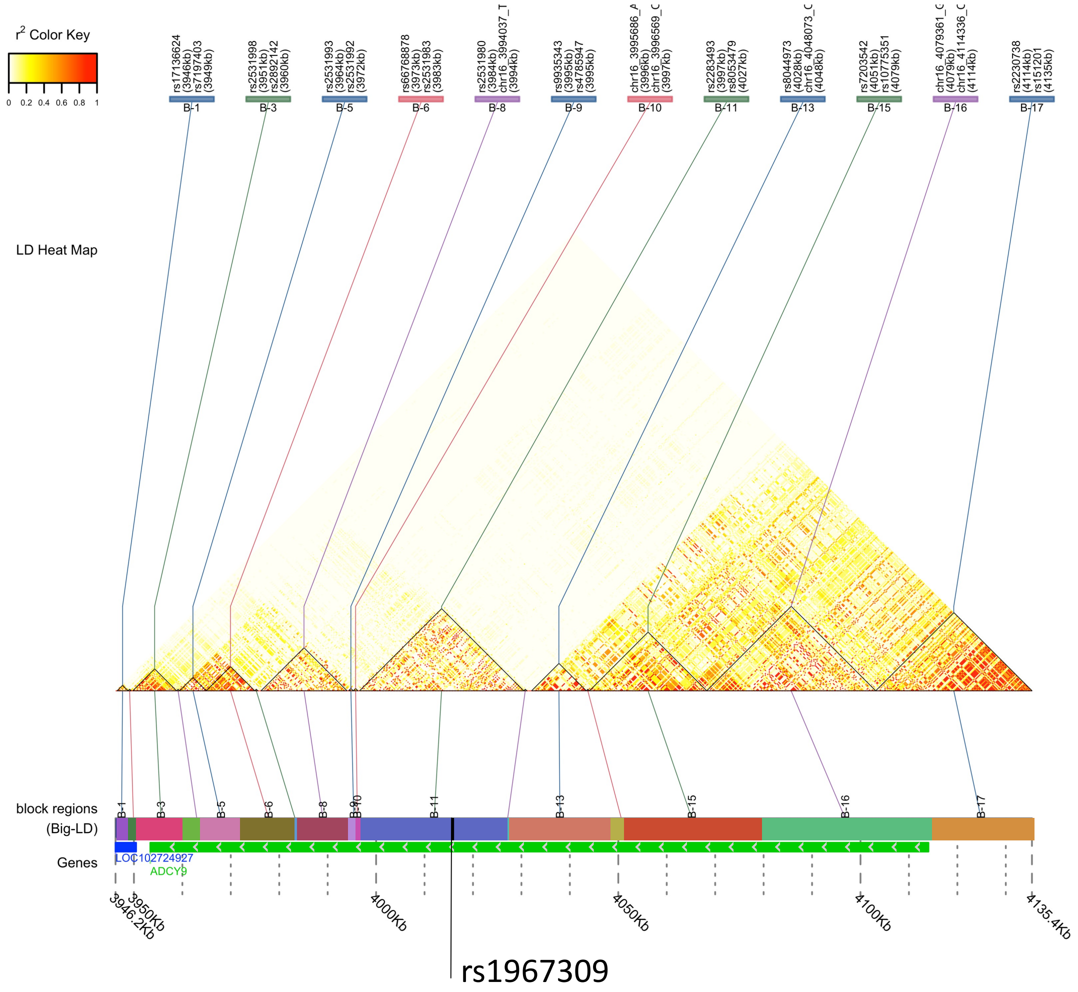


**Supplementary Figure 13. Gene structure of ADCY9 locus.**

Linkage Disequilibrium (LD) blocks within the ADCY9 locus, with surrounding gene identified below and grey arrows indicating 5' to 3'. Blocks were estimated in GTEx participants from European descent using BigLD of gpart package with CLQ cut at 0.3 for SNPs having a MAF above 5%. IDs of SNPs delimiting each LD block are indicated above. Position of the mutation rs1967309 is indicated in the corresponding LD block.

### Supplementary tables

**Supplementary Table 1.** Source and information about the continuous phenotypes used in the paper. The information included the source and the ID to retrieve the data, the consortium from which the data originated, the sample size, and whether the data is derived from a published paper or a laboratory source.

| Outcome | Source | Consortium | Sample Size | Origin |
|---|---|---|---|---|
| HDL cholesterol | ieu open gwas project : ieu-b-109 | UK Biobank | 403943 | PMID : 32203549 |
| LDL cholesterol | ieu open gwas project : ieu-b-110 | UK Biobank | 440546 | PMID : 32203549 |
| TG | ieu open gwas project : ieu-b-111 | UK Biobank | 441016 | PMID : 32203549 |
| Trunk fat-free mass | ieu open gwas project : ukb-b-17409 | MRC-IEU | 454508 | Ben Elsworth |
| Whole body fat-free mass | ieu open gwas project : ukb-b-13354 | MRC-IEU | 454850 | Ben Elsworth |
| Height | GWASatlas id : 3187 | UK Biobank | 385748 | PMID: 31427789 |
| Basal metabolic rate | ieu open gwas project : ukb-b-16446 | MRC-IEU | 454874 | PMID : NA |
| Thyroid Stimulating Hormone | ieu open gwas project : prot-a-530 | NA | 3301 | PMID : 29875488 |
| Birth weight | ieu open gwas project : ukb-b-13378 | MRC-IEU | 261932 | Ben Elsworth |
| Birth weight of first child | ieu open gwas project : ukb-b-3357 | MRC-IEU | 200272 | Ben Elsworth |
| FEV1 | ieu open gwas project : ukb-b-19657 | MRC-IEU | 421986 | Ben Elsworth |
| FVC | ieu open gwas project : ukb-b-7953 | MRC-IEU | 421986 | Ben Elsworth |

**Supplementary Table 2.** Source and information about the discrete phenotypes used in the paper. The information included the source and the ID required to retrieve the data, the consortium from which the data originated, the number of cases and controls included in the study, and the reference to the paper where the data is reported.

| **Outcome** | **Source** | **Consortium** | **Number of Cases** | **Number of Controls** | **PMID** |
|---|---|---|---|---|---|
| CVD-related | van der Harst, Pim (2017), “CAD meta-analysis”, Mendeley Data, V1, doi: 10.17632/gbbsrpx6bs.1 | UK Biobank, CARDIoGRAMplusC4D, Myocardial Infarction Genetics, CARDIoGRAM Exome | 122733 | 424528 | PMID : 29212778 |
| Early age-related macular degeneration | ieu open gwas project : ebi-a-GCST010723 | International AMD genomics consortium (IAMDGC) | 14034 | 91214 | PMID : 32843070 |
| Hypothyroidism | GWASatlas id : 4376 | NA | 3440 | 49983 | PMID : 30367059 |
| Hyperthyroidism | GWASatlas id : 4375 | NA | 1840 | 49983 | PMID : 30367059 |
| Miscarriage stillbirth | pheweb.org/UKB-TOPMed : 634 | UK Biobank | 5214 | 212254 | PMID : NA |

**Supplementary files**

*Supplementary file 1.* List of the first underlying cause of death (DTHFUCOD) from GTEx dataset, specifically belonging to the variables MHHRTATT and MHHRTDIS. These variables provide information about the CAD-related causes of death for the individuals included in the GTEx dataset.

*Supplementary file 2.* List of the first underlying cause of death (DTHFUCOD) from GTEx dataset that were removed from the control group in our analysis. This list includes the cause of death associated to the variable MHHRTDISB, the cause of death associated to heart but not linked to MHHRTATT and MHHRTDIS, as well as cases where the cause of death is unknown.

*Supplementary file 3.* Statistical summary of all SNPs that were used in at least one Mendelian Randomisation analysis conducted on thyroid tissue for the three exposures: eQTL of gene-level *CETP* expression, sQTL of Alternative splicing of exon 9 (AS9) and sQTL of alternative exon 1 (AS1). The information presented includes the beta coefficient, the effect allele, the p-value (pval) of the association, the standard error (se), the derived F-statistic (Fstat) of each SNP for the three exposures. We also identified the outcomes for which the SNP was used and in which analysis. The SNPs included in this list are those that passed the filtering criteria, which required a derived F-statistic of at least 10, a p-value below 0.001, and a minor allele frequency above 0.01 for at least one of the three exposures. Additionally, the SNPs were filtered based on correlation (r2>0.90) before being included in the analysis.

*Supplementary file 4.* Genetic correlation matrix for all SNPs that were used in at least one Mendelian Randomisation analysis. The correlation matrix was calculated using the ld_matrix function from the R package ieugwasr and was performed specifically on the 699 individuals of European descent in the GTEx dataset. The SNPs included in this matric are those that passed the filtering criteria, which required a derived F-statistic of at least 10, a p-value below 0.001, and a minor allele frequency above 0.01 for at least one of the three exposures. Additionally, the SNPs were filtered based on correlation (r2>0.90) before being included in the analysis.

### References


1. Robinson JT, Thorvaldsdóttir H, Winckler W, Guttman M, Lander ES, Getz G, et al. Integrative Genomics Viewer. Nat Biotechnol. 2011 Jan;29(1):24–6.
2. Lappalainen T, Sammeth M, Friedländer MR, ‘t Hoen PAC, Monlong J, Rivas MA, et al. Transcriptome and genome sequencing uncovers functional variation in humans. Nature. 2013 Sep;501(7468):506–11.
3. Alfieri C, Birkenbach M, Kieff E. Early events in Epstein-Barr virus infection of human B lymphocytes. Virology. 1991 Apr;181(2):595–608.
4. Jiang S, Zhou H, Liang J, Gerdt C, Wang C, Ke L, et al. The Epstein-Barr Virus Regulome in Lymphoblastoid Cells. Cell Host Microbe. 2017 Oct 11;22(4):561-573.e4.
5. Wang LW, Wang Z, Ersing I, Nobre L, Guo R, Jiang S, et al. Epstein-Barr virus subverts mevalonate and fatty acid pathways to promote infected B-cell proliferation and survival. PLOS Pathogens. 2019 Sep 13;15(9):e1008030.
6. Apostolou F, Gazi IF, Lagos K, Tellis CC, Tselepis AD, Liberopoulos EN, et al. Acute infection with Epstein–Barr virus is associated with atherogenic lipid changes. Atherosclerosis. 2010 Oct 1;212(2):607–13.
7. Mizuno A, Okada Y. Biological characterization of expression quantitative trait loci (eQTLs) showing tissue-specific opposite directional effects. Eur J Hum Genet. 2019 Nov;27(11):1745–56.
8. Oliveira HC, Chouinard RA, Agellon LB, Bruce C, Ma L, Walsh A, et al. Human cholesteryl ester transfer protein gene proximal promoter contains dietary cholesterol positive responsive elements and mediates expression in small intestine and periphery while predominant liver and spleen expression is controlled by 5’-distal sequences. Cis-acting sequences mapped in transgenic mice. J Biol Chem. 1996 Dec 13;271(50):31831–8.
9. Schwarzer G, Carpenter JR, Rücker G. Meta-Analysis with R [Internet]. Cham: Springer International Publishing; 2015 [cited 2023 May 29]. (Use R!). Available from: https://link.springer.com/10.1007/978-3-319-21416-0
10. Duell PB, Bierman EL. The relationship between sex hormones and high-density lipoprotein cholesterol levels in healthy adult men. Arch Intern Med. 1990 Nov;150(11):2317–20.
11. Phillips GB, Pinkernell BH, Jing TY. Relationship Between Serum Sex Hormones and Coronary Artery Disease in Postmenopausal Women. Arteriosclerosis, Thrombosis, and Vascular Biology. 1997 Apr;17(4):695–701.
12. Anagnostopoulou KK, Kolovou GD, Kostakou PM, Mihas C, Hatzigeorgiou G, Marvaki C, et al. Sex-associated effect of CETP and LPL polymorphisms on postprandial lipids in familial hypercholesterolaemia. Lipids Health Dis. 2009 Jun 26;8:24.
13. Lim GB. Role of sex hormones in cardiovascular diseases. Nat Rev Cardiol. 2021 Jun;18(6):385–385.
14. Poulin SP, Dautoff R, Morris JC, Barrett LF, Dickerson BC. Amygdala atrophy is prominent in early Alzheimer’s disease and relates to symptom severity. Psychiatry Res. 2011

Oct 31;194(1):7–13.
15. Arias-Vásquez A, Isaacs A, Aulchenko YS, Hofman A, Oostra BA, Breteler M, et al. The cholesteryl ester transfer protein (CETP) gene and the risk of Alzheimer's disease. Neurogenetics. 2007 Aug;8(3):189–93.
16. Murphy EA, Roddey JC, McEvoy LK, Holland D, Hagler DJ, Dale AM, et al. CETP polymorphisms associate with brain structure, atrophy rate, and Alzheimer's disease risk in an APOE-dependent manner. Brain Imaging Behav. 2012 Mar;6(1):16–26.
17. Tawakol A, Ishai A, Takx RA, Figueroa AL, Ali A, Kaiser Y, et al. Relation between resting amygdalar activity and cardiovascular events: a longitudinal and cohort study. The Lancet. 2017 Feb 25;389(10071):834–45.
18. Gianaros PJ, Hariri AR, Sheu LK, Muldoon MF, Sutton-Tyrrell K, Manuck SB. Preclinical Atherosclerosis Covaries with Individual Differences in Reactivity and Functional Connectivity of the Amygdala. Biol Psychiatry. 2009 Jun 1;65(11):943–50.
19. Hou H, Ma R, Guo H, He J, Hu Y, Mu L, et al. Association between Six CETP Polymorphisms and Metabolic Syndrome in Uyghur Adults from Xinjiang, China. Int J Environ Res Public Health. 2017 Jun;14(6):653.
20. Guo S, Hu Y, Ding Y, Liu J, Zhang M, Ma R, et al. Association between Eight Functional Polymorphisms and Haplotypes in the Cholesterol Ester Transfer Protein (CETP) Gene and Dyslipidemia in National Minority Adults in the Far West Region of China. Int J Environ Res Public Health. 2015 Dec;12(12):15979–92.
21. Quanjer PH, Hall GL, Stanojevic S, Cole TJ, Stocks J. Age- and height-based prediction bias in spirometry reference equations. European Respiratory Journal. 2012 Jul 1;40(1):190–7.
22. Broere-Brown ZA, Baan E, Schalekamp-Timmermans S, Verburg BO, Jaddoe VWV, Steegers EAP. Sex-specific differences in fetal and infant growth patterns: a prospective population-based cohort study. Biology of Sex Differences. 2016 Dec 3;7(1):65.
23. Vaquero-Garcia J, Barrera A, Gazzara MR, González-Vallinas J, Lahens NF, Hogenesch JB, et al. A new view of transcriptome complexity and regulation through the lens of local splicing variations. Valcárcel J, editor. eLife. 2016 Feb 1;5:e11752.
